# Diversity and habitat-associated distribution of bacterial and protistan communities in *Phragmites australis* biofilms and surrounding waters of Masurian lakes (Poland)

**DOI:** 10.1101/2025.11.26.690845

**Authors:** Valentina Smacchia, Michał Karlicki, Bartosz Kiersztyn, Anna Karnkowska

## Abstract

Epiphytic biofilms on aquatic macrophytes represent distinct microbial habitats within freshwater ecosystems, yet how individual taxa are distributed between biofilm and planktonic communities remains poorly characterised, particularly when bacteria and protists are examined simultaneously. Here we characterise bacterial and protistan communities associated with *Phragmites australis* stem biofilms and surrounding waters across five lakes in the Great Masurian Lake District, northeastern Poland, using 16S and 18S rRNA gene metabarcoding. In total, we identified 5,332 bacterial and 4,509 protistan amplicon sequence variants (ASVs). Bacterial biofilm samples harboured higher ASV richness than water samples across all five lakes, with a proportionally larger rare and spatially restricted fraction, while planktonic communities were dominated by abundant, widely distributed ASVs. Despite this extensive rare fraction, only 43 bacterial and 7 protistan ASVs were consistently detected across all biofilm samples while remaining absent from the water column. A widespread component was shared between habitats, though taxa abundant in water tended to occur at lower abundance in the biofilm. Indicator value analysis and Random Forest classification identified taxa characteristic of each habitat, with bacterial communities showing stronger habitat associations and greater cross-lake consistency than protist assemblages, whose quantitative composition varied considerably across lakes regardless of habitat. These findings demonstrate that occurrence and abundance capture complementary and sometimes contrasting dimensions of habitat specificity, as rare taxa dominate biofilm richness yet contribute little to its overall composition. Moreover, habitat structuring operates differently for bacteria and protists, underlining the value of examining multiple microbial domains simultaneously.

## Introduction

Microbial communities in freshwater ecosystems are complex and dynamic, occurring in both free-living planktonic assemblages and surface-associated communities such as biofilms. Biofilms are surface-attached microbial communities embedded in an extracellular matrix that form distinct microhabitats with physical, chemical, and biological conditions that can differ from those of the surrounding water [1–3].

Aquatic plants provide important surfaces for the development of epiphytic biofilms, which represent a major component of freshwater microbial communities [4]. The plant surface provides a heterogeneous substrate for microbial colonisation, supporting complex communities. In lake ecosystems, epiphytic bacterial communities associated with macrophytes have been shown to be diverse and taxonomically distinct among plant-associated habitats [5].

One of the most widespread emergent macrophytes in temperate freshwater ecosystems is *Phragmites australis*, which frequently forms extensive stands along lake margins and other shallow water bodies. Its stems support diverse epiphytic microbial communities, which have been investigated across different freshwater systems [6–8]. Previous studies in lakes have shown that bacterial communities developing on *P. australis* stems can differ markedly from those in the surrounding lake water, and that these communities may change during biofilm development and across seasons [7, 8]. Studies of reed-associated biofilms have also documented diverse bacterial assemblages and differences in their physiological profiles [6, 9].

Biofilm communities are connected to the surrounding planktonic microbial community and may share a proportion of their microbial taxa while maintaining distinct community structures. For example, in interconnected shallow ponds, biofilm and bacterioplankton communities were highly distinct, although both were structured by similar environmental processes [10]. In Lake Baikal, biofilm and planktonic bacterial communities showed clear differences in their overall structure, while sharing several major bacterial groups [11]. A study in Lake Erie likewise identified both biofilm- and plankton-associated taxa, with only a relatively small fraction of taxa shared between the habitats [12]. Thus, the relationship between these habitats is characterised by both taxonomic overlap and community differentiation: biofilms are connected to the surrounding water, but the presence of shared taxa does not imply equivalent community composition. An important remaining question is therefore not whether biofilm and planktonic communities differ, but how individual taxa are distributed between these habitats. Despite growing evidence for compositional differences between the two, relatively little is known about how specific taxa are distributed across biofilm and planktonic habitats in freshwater lakes. In particular, few studies have simultaneously examined both bacterial and protistan communities in this context.

To address this gap, we analysed bacterial and protist communities associated with *P. australis* stem biofilms and compared them with planktonic communities in five lakes of the Great Masurian Lake District (GMLD), northeastern Poland. The GMLD comprises dimictic temperate lakes representing a range of trophic states [13]. Using 16S and 18S rDNA metabarcoding we aimed to: 1) characterise bacterial and protistan community composition and diversity across biofilm and planktonic habitats; (2) examine the distribution of individual ASVs across habitats, considering their occurrence, relative abundance, and consistency across lakes; and (3) identify taxa showing characteristic associations with biofilm and planktonic habitats. Community-level physiological profiles were additionally examined as exploratory context for the compositional findings.

## Experimental Procedures

### Sampling

Five lakes in the Great Masurian Lake District (GMLD, NE Poland) were selected as sampling sites (Fig. 1): Ryńskie (RYN), Roś (ROS), Mikołajskie (MIK), Przystań (PRZ) and Majcz (MAJ). First four of them are part of the Great Masurian Lake System (GMLS). The selected lakes share broadly similar geological, climatic and catchment characteristics, reflecting the regional context, but differ in their trophic conditions and levels of anthropogenic pressure (Table S1, Fig. 1). Their distribution across the region and the environmental variation among them provided an opportunity to explore microbial communities across a range of conditions characteristic of Masurian lakes.

**Fig. 1.**
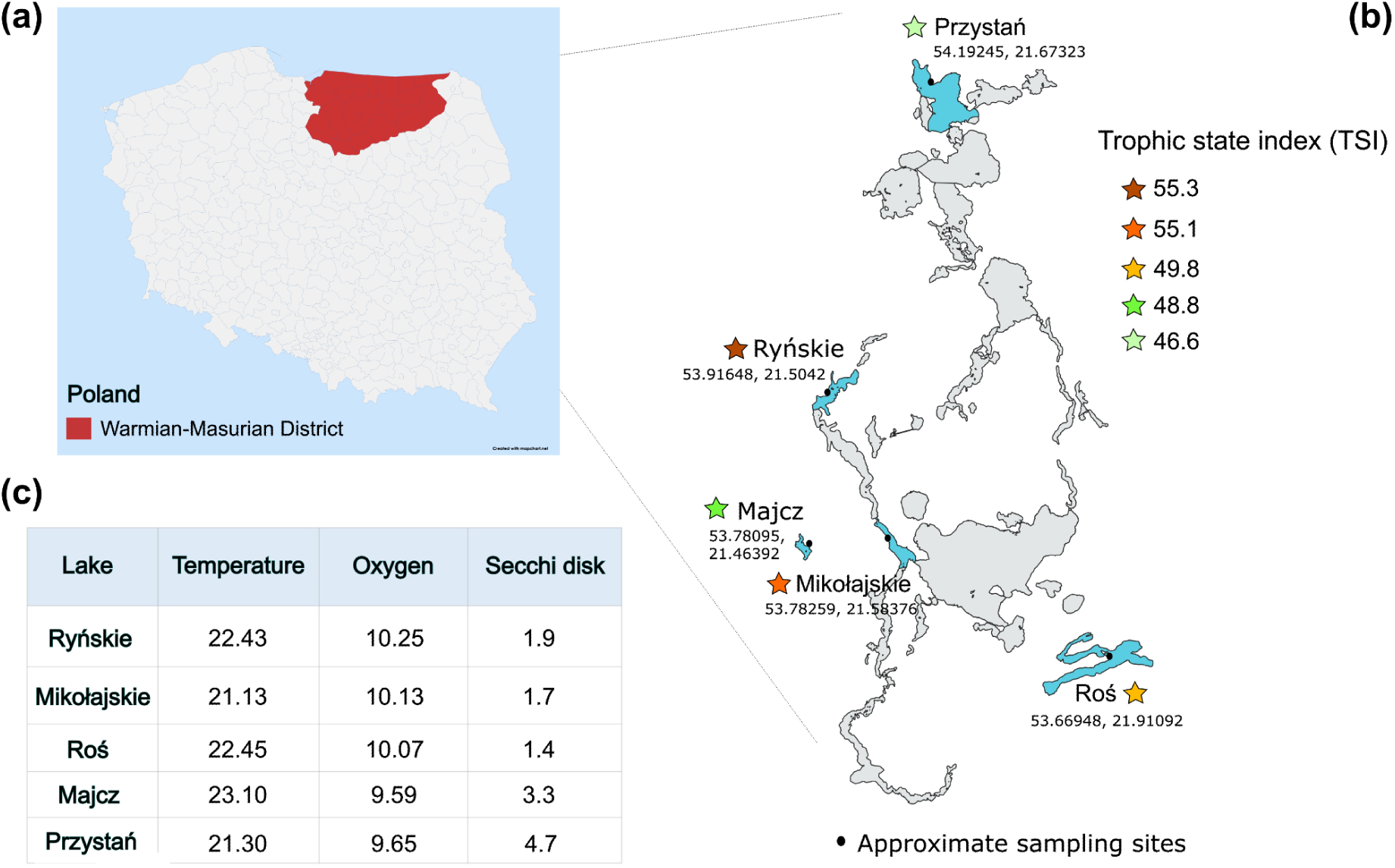
Geographic location of the studied lakes. (**a**) The map shows the Warmia-Masurian District in northeastern Poland (highlighted in red, upper left inset), where the Great Masurian Lake District (GMLD) is located. (**b**) The five sampled lakes are shown with marked sampling sites. Lakes are classified according to their Trophic State Index (TSI). (**c**) Key physicochemical parameters — water temperature (°C), dissolved oxygen (mg/L), and water transparency (Secchi disk depth, m) — are reported for each lake.

Sampling was conducted between 25 and 27 July 2022, with the exact sampling date for each lake reported in Table S1. From each lake, four subsamples were collected: biofilm from *Phragmites australis* stems (B), littoral water within (LP) and outside (L) the *P. australis* field, and pelagic water (P). L samples were collected at least 10 m from the nearest *P. australis* field, and P samples at least 200 m from the littoral zone. The three water-sampling zones were selected to represent a gradient of increasing distance from the reed stands, allowing us to explore potential variation in microbial community composition between water closely associated with *P. australis* vegetation, adjacent littoral water, and more distant pelagic waters. At each site, 5 L of surface water was collected. Biofilm samples from *P. australis* stems were collected following Tsuchiya et al. (2011) with modifications. Stems were cut into 30 cm sections, starting 5 cm below the water surface and extending to a depth of 35 cm; the upper 10 cm were discarded, and ten fragments per lake were retained and placed in a sterile plastic bottle filled with lake water from the corresponding *P. australis* field. Biofilms from 10 cm segments were removed using a sterile brush and suspended in 100 ml of 0.2 µm-filtered site water. For taxonomic composition analysis, each water and biofilm sample was filtered under low air pressure using Nucleopore filters (Whatman, Maidstone, UK) with a 0.2 μm pore size. The filters were stored at −80 °C until DNA extraction.

### Physical and chemical metadata

In situ temperature and oxygen concentration were measured at each sampling point using a YSI ProODO probe (Yellow Springs, YSI Inc./Xylem Inc., USA). Photic zone depth was estimated using a LiCor LI-250A with LI-193R PAR sensor and a Secchi disk. The Trophic State Index (TSI) was calculated as the mean of values derived from total phosphorus measured over three consecutive summers (2021–2023; Table S1) using the Carlson formula [14]. Dissolved organic carbon (DOC), total phosphorus (TP), phosphate (PO₄³⁻), and total nitrogen (TN) concentrations were determined as described in Kiersztyn et al., (2019).

### Physiological fingerprint diversity and Leucine-aminopeptidase activity

The Biolog® EcoPlates™ (Biolog Inc., Hayward, CA, USA) method was used to determine community-level physiological profiles, with modifications as described by Kiersztyn et al., (2019). Maximal potential leucine-aminopeptidase activity (Vmax_LAP) was measured fluorometrically following the methods of Chróst and Siuda (2006) and Kiersztyn et al. (2012). For both measurements, a volume correction factor was applied to allow approximate comparison between biofilm suspensions and water samples, calculated as the ratio of the suspension volume (100 ml) to the mean volume of a 10 cm stem segment; results were expressed per sample volume and were not normalised to biomass or cell abundance. These data are provided in supplementary material (Tables S2 and S3, Fig. S1).

### DNA extraction, amplification, and sequencing

DNA was isolated using the GeneMATRIX Soil DNA Purification Kit (EURx Sp. z o.o., Gdańsk, PL) from filters, following the manufacturer’s protocol with minor modifications: after the addition of 60 µL lysis buffer, samples were homogenised for 10 minutes with a vertical homogeniser, stored at −80℃ for 1 hour, and homogenised again for 5 minutes at maximum speed. For the final elution, 30 µL DNase/RNase-free water was used.

Extracted DNA was quantified using a NanoDrop (Implen/Thermo Fisher Scientific, USA), and fragment size was verified on a 1% agarose gel. Samples were sequenced at the University of Warsaw (CeNT Genomics Core Facility, Poland) on a NovaSeq 6000 platform following amplification and library preparation.

The prokaryotic V4-V5 and eukaryotic V4 hypervariable regions of the 16S and 18S rRNA genes, respectively, were amplified following [18, 19] with minor modifications: primer concentrations were increased to 0.2 µM for prokaryotes and reduced to 0.4 µM for eukaryotes; for the prokaryotic primers, 20 cycles were used for amplification. Amplifications were performed in triplicate, pooled, and purified using a PCR clean-up kit (Syngen).

### Sequence processing and data analysis

Sequence quality checks were performed on raw sequencing data using FastQC [20]. Reads were imported into the QIIME2 environment for demultiplexing, adapter trimming, and primer removal [21, 22]. Denoising was conducted using the DADA2 algorithm in QIIME2 [23].

Taxonomic assignment of the V4-V5 16S rRNA and V4 18S rRNA sequences was performed using USEARCH global alignment implemented in VSEARCH [24]. A minimum query coverage of 90% and a minimum identity threshold of 70% were applied to all sequences. The full taxonomy of the best hit was assigned to each ASV, with the 70% identity threshold serving as a minimum cutoff to exclude clearly unrelated sequences. Same parameters were applied to both markers to ensure methodological consistency between prokaryotic and eukaryotic community analyses.

The SILVA 138.1 database [25] was used for 16S rRNA classification, while the Protist Ribosomal Database PR2 5.0.0 was used for 18S rRNA classification [26]. All taxonomic and diversity analyses were conducted in RStudio IDE using the phyloseq package [27, 28] after removal of ASVs classified as Eukaryota, chloroplasts, or mitochondria from the 16S dataset, and ASVs assigned to Metazoa, Embryophyta, Bacteria, or Archaea from the 18S dataset.

The analyses, packages used, and the analytical workflow were based on the code published by Karlicki et al. (2024) with modifications (Zenodo DOI 10.5281/zenodo.17552625). Eukaryotic and prokaryotic prevalence analyses were performed using the microbiome and microViz packages [30, 31]. Taxonomic analyses were conducted on non-normalised data, transformed to relative abundances of ASVs. For subsequent analyses, samples were grouped by zones: biofilm, littoral inside and outside the Phragmites australis field, and the pelagic zone. To visualise the distribution of samples in ordinated space, a Principal Coordinates Analysis (PCoA) based on Bray-Curtis dissimilarity was performed; this analysis was used for descriptive purposes only given the absence of replicates. Species richness was estimated using the iNEXT package [32], applying sample-size-based rarefaction and extrapolation curves (curve type 3) to visualise and compare richness across individual samples. Hill numbers (q = 0, 1, and 2) were also calculated as exact diversity indices for each sample (Table S4). The distribution of ASVs across habitats was examined by classifying them according to their occupancy (the proportion of samples within each zone in which they occurred) and mean relative abundance, calculated across all samples of each zone. A threshold of 0.1% relative abundance was selected as an operational criterion to distinguish relatively abundant from low-abundance ASVs, considering both thresholds previously applied in microbial community studies [33–35] and the distribution of relative abundances observed in our dataset. Based on these criteria, four categories were defined: “abundant and ubiquitous,” for ASVs present in all samples of a given zone with a mean relative abundance ≥0.1%; “rare and ubiquitous,” for ASVs present in all samples but with a mean relative abundance <0.1%; “abundant and patchy,” for ASVs with a mean relative abundance ≥0.1% but not present in all samples; and “Rare and patchy,” comprising the remaining ASVs with a mean relative abundance <0.1% that were not present in every sample.

### Consistency across lakes

To assess whether ubiquitous ASVs also showed consistent relative abundances across lakes, lake-to-lake variability was evaluated separately for each zone and domain. The analysis was restricted to ubiquitous ASVs, defined as ASVs detected in all five lakes within a given zone, and abundant and rare ubiquitous ASVs were retained as separate categories according to the abundance threshold defined above. For each ASV, the coefficient of variation (CV) of relative abundance across the five lakes was calculated as the standard deviation divided by the mean relative abundance. For descriptive purposes, ASVs were grouped into low (CV < 0.5), intermediate (0.5 ≤ CV ≤ 1), and high (CV > 1) lake-to-lake variability.

Because overall variability among lakes does not necessarily imply dominance by a single lake, the concentration of each ASV’s abundance in individual lakes was evaluated separately. For each ubiquitous ASV, we calculated the proportion of its cumulative relative abundance across the five lakes contributed by the lake in which it reached its highest relative abundance. A contribution >50% indicated that a single lake accounted for more of the ASV’s cumulative abundance than the other four lakes combined, while contributions >70% and >90% were used descriptively to identify increasingly pronounced cases of single-lake concentration (Table S5).

### Indicator species analysis and Random Forest

Indicator species analysis was performed separately for bacteria and protists using the multipatt function of the R package *indicspecies* [36] to identify ASVs associated with individual zones or combinations of zones. All possible combinations of zones were considered (duleg = FALSE), and the permutation procedure was based on 9,999 permutations. Indicator values were decomposed into specificity (A) and fidelity (B). A measures the strength of the association of an ASV with the target zone or zone combination, whereas B represents its frequency of occurrence within the target group. To focus on taxa showing both strong habitat specificity and consistent occurrence, characteristic ASVs were defined using effect-size criteria of A ≥ 0.8 and B = 1, rather than on statistical significance alone, as the small number of biofilm samples limits the resolution of permutation-based p-values and their ability to detect strong associations. This conservative selection was intended to distinguish consistently occurring and strongly associated taxa from ASVs showing weaker or less recurrent associations.

To identify 16S rDNA and 18S rDNA ASVs whose abundance correlated with the matrix group (biofilm: n = 5; water: n = 15 per sequencing marker) across samples, we applied a supervised machine learning algorithm, Random Forest (RF) [37], following the methodology of Karlicki et al. (2024). Initially, the three aquatic zones (L, LP and P) were treated as separate classes. However, preliminary RF models showed substantial misclassification among these zones, with an out-of-bag (OOB) error rate of ≥70%, indicating limited discrimination among aquatic communities. Therefore, the three aquatic zones were combined into a single water class, allowing the analysis to focus on differences between biofilm and aquatic assemblages. For each dataset, we selected the top 60 ASVs with the highest mean decrease in the Gini coefficient index (Fig. S2), representing those with the greatest impact on sample classification. The classification achieved an out-of-bag (OOB) error rate of 0% for the 18S rDNA dataset and 5% for the 16S rDNA dataset. Given the small sample size, the RF model was used as a feature-selection tool to identify ASVs that discriminate between biofilm and water assemblages, rather than as a predictive classifier; the OOB error rates are reported for completeness but should be interpreted with caution in this context.

## Results

### Community structure of biofilm and planktonic assemblages

Metabarcoding sequencing generated 2.96 million 16S rDNA and 3.35 million 18S rDNA reads, of which 43.84% and 62.15% passed quality filtering, yielding 5,332 bacterial and 4,509 eukaryotic ASVs, respectively (Table S6, Fig. S3). The most prevalent bacterial phyla in both biofilm and water samples were Proteobacteria (21–67%), Actinobacteriota (0.2–42.8%) and Bacteroidetes (15.2–41.4%) (Fig. 2a, left panel). The protist community mostly consisted of Dinoflagellata (3.4–89.4%), Gyrista (1.5–38%), Perkinsea (0.7–28%), Ciliophora (2.5–24%), Chlorophyta (0.9–25.7%), Fungi (0.6–39.2%) and Cercozoa (0.16–30.9%) (Fig. 2a, right panel). While the major phyla were similar among samples (Fig. 2a), differences emerged at lower taxonomic levels (Fig. S4), with bacterial biofilm communities appearing to contain more taxa per sample than planktonic ones as shown from iNEXT curves in Fig. 2b (Table S4, S6).

**Fig. 2.**
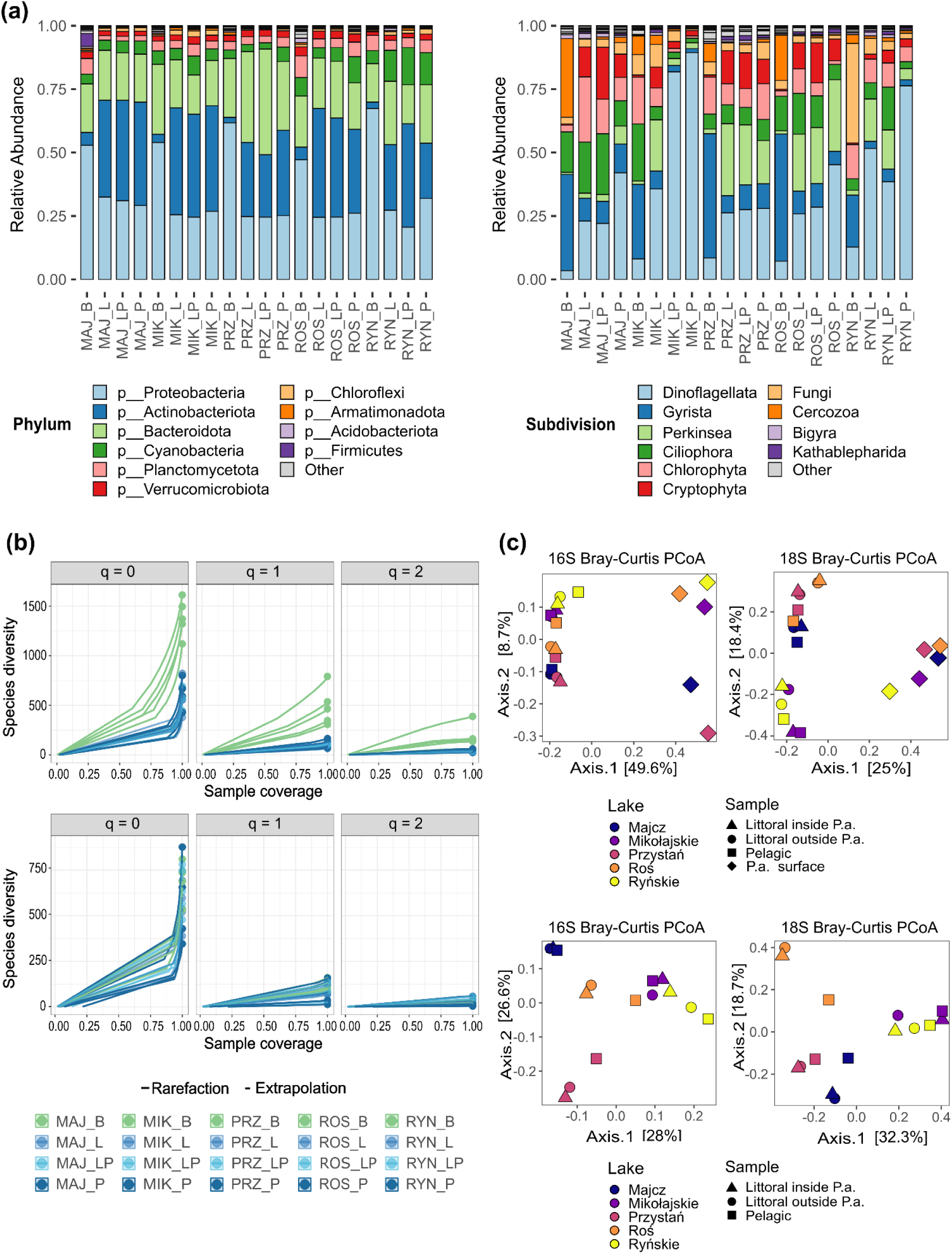
Microbial community composition and taxon distribution in biofilm and planktonic samples across five lakes. (**a**) Relative abundance of bacterial (left panel) and protist (right panel) communities at phylum and subdivision level, respectively, across all biofilm and water samples. Samples are grouped by lake (Majcz, Mikołajskie, Przystań, Roś, Ryńskie) and sample type (biofilm, littoral water inside and outside *P. australis* bed, pelagic water). (**b**) Sample-size-based rarefaction and extrapolation curves for bacterial (upper panels) and protist (lower panels) ASVs, shown for Hill numbers q = 0, q = 1, and q = 2 as a function of sample coverage. Colors indicate sample type, with green representing biofilm-associated samples and shades of blue representing water samples. Solid and dashed lines represent rarefaction and extrapolation, respectively, and points indicate observed values. (**c**) Principal Coordinate Analysis (PCoA) of Bray-Curtis dissimilarity for bacterial (16S, left) and protist (18S, right) communities. Upper panels show all sample types distinguished by shape, with lakes colour-coded; lower panels show only water samples to highlight between-lake variation. P.a. = *Phragmites australis.*

PCoA of Bray-Curtis dissimilarity showed that biofilm and planktonic communities occupied visually distinct regions of ordination space for both prokaryotes and protists, while samples from the three water zones broadly overlapped (Fig. 2c, upper panel). Given that each lake contributed a single biofilm sample, formal statistical comparisons between habitat types were not pursued, as lake identity and habitat effects cannot be fully disentangled. The visual separation between biofilm and water communities was nonetheless consistent across all five lakes and both taxonomic domains. Within the water samples, those from the three aquatic zones of the same lake tended to cluster together, while samples from different lakes appeared more separated in the ordination (Fig. 2c, lower panel), suggesting that between-lake variation outweighs compositional differences among water zones within lakes. The distance gradient from the reed stands may therefore not represent a major structuring axis at the scale sampled, and the three water zones are treated collectively in analyses where zone-level resolution does not add descriptive value.

Coverage-based rarefaction and extrapolation with Hill numbers (iNEXT) were used to assess and compare the diversity of bacterial and protist ASVs across samples (Fig. 2b). Bacterial biofilm samples were characterized by higher ASV richness (q0 = 1,120–1,613), than water samples (q0 = 378–823), with Hill numbers decreasing at q1 (306.6–791.1 vs. 63.7–163.8) and q2 (56.4–387.7 vs. 22.5–60.2) in both sample types (Fig 2b and Table S4). In contrast, protist richness showed broadly overlapping ranges between biofilm and water samples (q0 = 533–811 and 346–877, respectively), with diversity decreasing from q0 to q1 and q2 in both types of samples (Fig 2b and Table S4). Together, these patterns indicate high bacterial richness (q0) in biofilm, while the patterns in effective diversity (q1, q2) were less pronounced, suggesting a larger proportion of rare bacterial taxa in the biofilm. Protist richness and diversity showed more similar ranges across habitat types at all diversity orders.

### Occupancy–abundance patterns across biofilm and planktonic habitats

Across all zones and domains, the proportion of abundant ASVs increased with occupancy: abundant taxa were nearly absent at low occupancy (ASVs present in only one or two samples), whereas their proportion increased progressively towards full occupancy (Fig. 3, Table S7). This pattern was particularly pronounced for bacteria, where abundant ASVs remained low at intermediate occupancy levels (4.7–5.8% at 3/5 samples and 13.8–24.0% at 4/5 samples across zones) before increasing sharply among fully ubiquitous ASVs (46.0–71.3%).

**Fig. 3.**
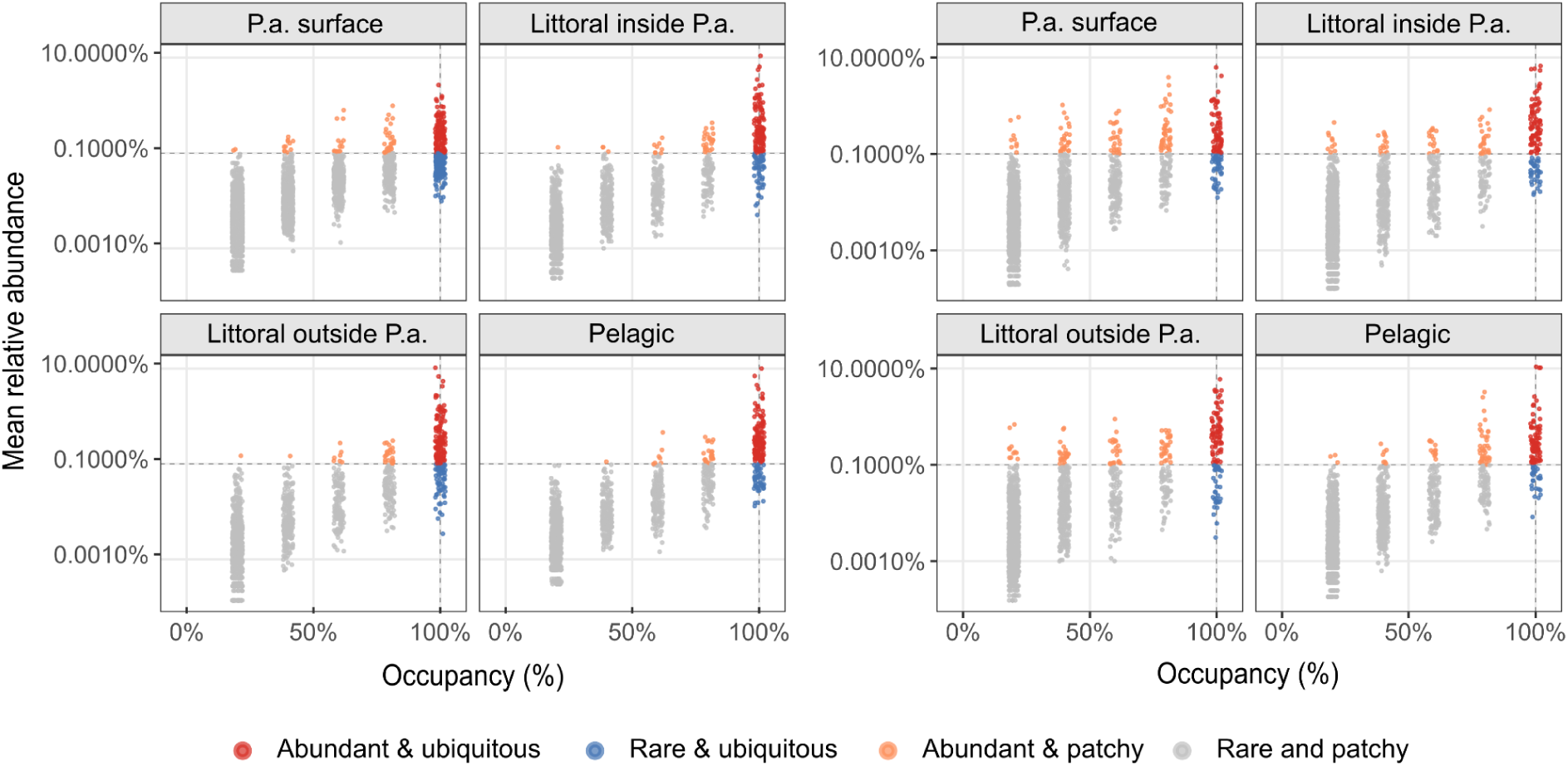
Occupancy–abundance relationships of bacterial and protist ASVs across sampling zones. Occupancy–abundance relationships for bacterial (left) and protist (right) ASVs in biofilm (Phragmites australis), littoral water inside and outside the P. australis bed, and pelagic water. Occupancy represents the fraction of samples within each zone in which an ASV was detected, while the y-axis shows its mean relative abundance. ASVs are classified according to their occupancy and abundance patterns as abundant and ubiquitous, abundant and patchy, rare and ubiquitous, or rare and patchy. Dashed lines indicate the thresholds used for classification.

Protists showed the same overall relationship, although with a different pattern in the biofilm. In the water zones, the proportion of abundant ASVs increased markedly from 4/5 occupancy (28.2–35.8%) to full occupancy (68.0–75.7%), broadly resembling the bacterial pattern (Table S7). In the biofilm, the increase was more gradual, with abundant ASVs already representing 38.7% of those occurring in 4/5 samples and 59.6% of fully ubiquitous ASVs.

These patterns were also reflected in the contribution of ubiquitous ASVs to total relative abundance. In the water zones, ubiquitous ASVs accounted for 81.4–83.7% of bacterial and 62.3–68.9% of protist relative abundance, largely due to the abundant and ubiquitous fraction (78.9–80.7% and 60.8–67.3%, respectively), whereas rare ubiquitous taxa contributed only 2.5–3.1% and 1.3–1.6% (Table S7). In the biofilm, by contrast, the contribution of ubiquitous ASVs decreased to 47.2% for bacteria and 39.7% for protists, but the remaining abundance was distributed differently between domains. In protists, abundant but patchy ASVs accounted for 39.2% of total relative abundance, nearly matching the contribution of the ubiquitous fraction itself. In bacteria, abundant but patchy ASVs contributed only 12.2%, whereas a larger proportion of total abundance was represented by the low-abundance, non-ubiquitous “Rare and patchy” category (40.5%).

Together, these patterns indicate that abundant taxa in the water zones were largely concentrated among ubiquitous ASVs, while in the biofilm, abundant taxa were more broadly distributed across occupancy levels, with this contrast being more pronounced for protists than for bacteria.

### Distribution across habitats

For both bacteria and protists, ASV richness and spatial distribution varied between the biofilm and water habitats. This contrast was particularly pronounced for bacteria: of the 5,332 bacterial ASVs detected, 3,945 occurred in the biofilm, compared with 2,375 across the water habitat as a whole and approximately 1,400 in each water zone (Table S7). Accordingly, most bacterial ASVs restricted to a single zone or habitat were associated with the biofilm, with 2,957 ASVs unique to the *Phragmites australis* surface compared with only 270–337 in each water zone (Table S7). For protists, overall richness was more evenly distributed between habitats: 2,072 of the 4,509 ASVs occurred in the biofilm and 3,244 across the water habitat, with 1,562–1,983 ASVs in the individual water zones (Table S7). Nevertheless, biofilm samples again contained the largest number of ASVs consistently detected in this habitat but not detected in the sampled water communities (1,265), compared with 443–650 ASVs detected exclusively in each water zone (Table S7).

Despite the large pool of ASVs unique to the biofilm, only a small subset was consistently detected across all biofilm samples. Among bacteria, only 43 ASVs were both ubiquitous within the biofilm and absent from all water zones (Fig. 4a and 4b, Table S7). Similarly, only 7 protist ASVs showed the same distribution pattern. In contrast, none of the ubiquitous ASVs in the individual water zones was truly restricted to that zone in either domain (Fig. 4b, Table S7). ASVs that were ubiquitous within a given water zone but not ubiquitous in the other zones were nevertheless detected elsewhere at lower occupancy (Fig. 4b). Thus, ubiquity within a zone did not necessarily imply exclusive occurrence within a sampled habitat or zone: while the biofilm contained a small set of ASVs consistently detected across biofilm samples but not detected in the sampled water communities, widespread ASVs in individual water zones were always part of a community extending beyond the individual water zone. This ubiquitous bacterial component detected exclusively in the sampled biofilm communities was taxonomically diverse and included ASVs assigned to genera such as *Flavobacterium* and *Phaeodactylibacter* (Bacteroidota), *Gemmatimonas* (Gemmatimonadota), *Paludibaculum* (Acidobacteriota), and *Ideonella*, *Rhodobacter* and *Sphingomonas* (Proteobacteria), among others. The corresponding protist component comprised seven ASVs, including *Encyonopsis* (Stramenopiles), *Tetracystis* (Chlorophyta), Salpingoecidae Group C2 and Cladochytriales (Opisthokonta), Pleurostomatida and Hypotrichia (Alveolata), and *Angulamoeba* (Evosea) (Table S8). Notably, ASVs consistently detected in biofilm samples but not detected in the sampled water communities did not necessarily belong to taxa restricted to biofilms, as many of these taxa were also represented by distinct ASVs detected in the water compartments.

**Fig. 4.**
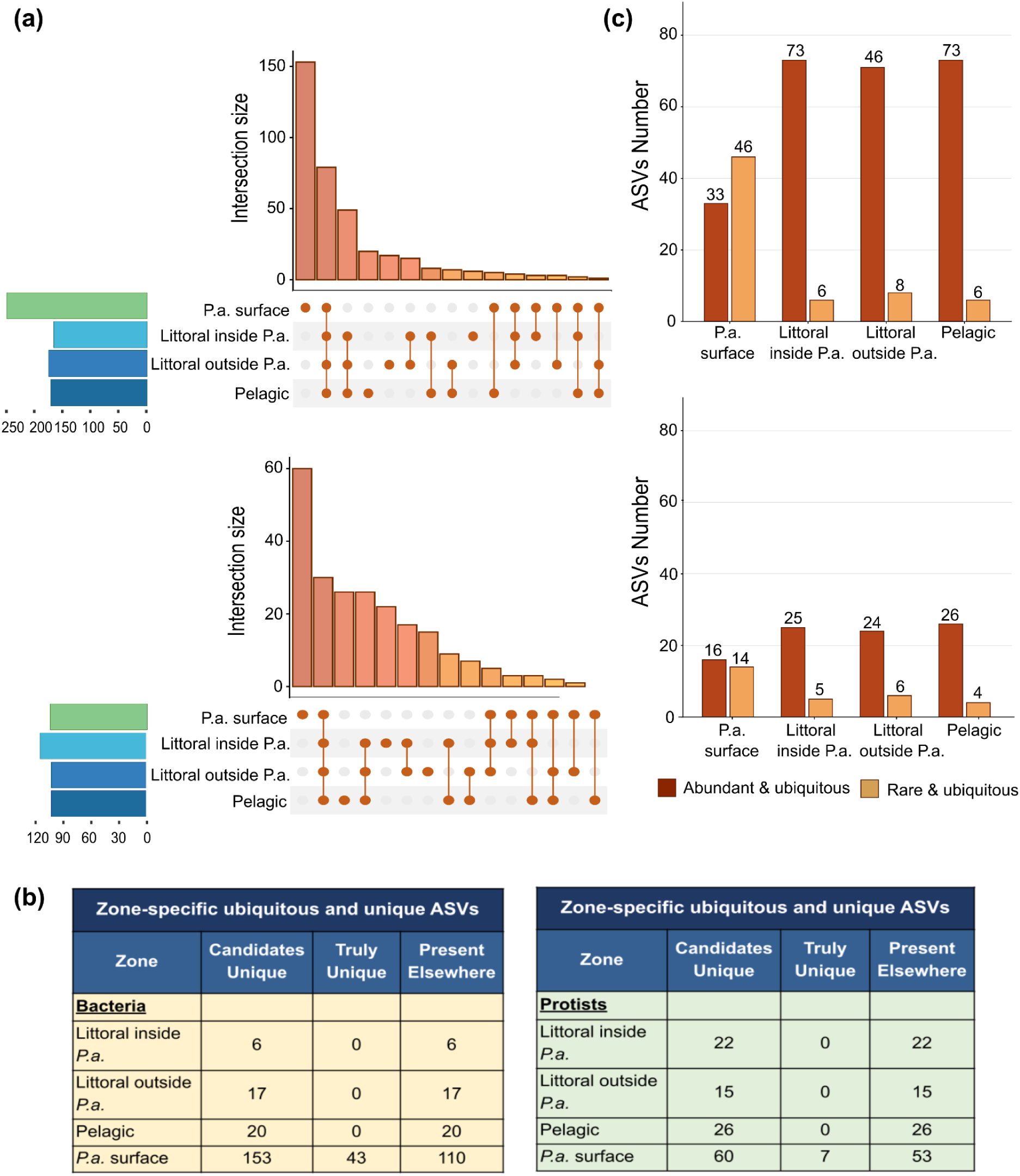
Distribution and occupancy–abundance classification of bacterial and protist ASVs across sampling zones. (**a**) UpSet plots showing the number of bacterial (upper panel) and protist (lower panel) ASVs shared among or restricted to biofilm (Phragmites australis surface), littoral water inside and outside the P. australis bed, and pelagic water. (**b**) Number of zone-specific candidate ubiquitous ASVs identified in each sampling zone, distinguishing ASVs exclusively occurring in that zone (“truly unique”) from those also detected in other zones. (**c**) Number of bacterial (upper panel) and protist (lower panel) ASVs classified as abundant and ubiquitous or rare and ubiquitous within each sampling zone according to the occupancy–abundance analysis.

The widespread component was also shared across habitats. Among bacteria, 128 ASVs were ubiquitous in all three water zones, of which 79 were also ubiquitous in the biofilm (Fig. 4a). An additional 18 bacterial ASVs ubiquitous in the biofilm were also ubiquitous in one or two, but not all three, water zones. Similarly, 56 protist ASVs were ubiquitous across all three water zones, 30 of which were also ubiquitous in the biofilm, while a further 14 biofilm-ubiquitous ASVs were ubiquitous in one or two water zones (Fig. 4a). The remaining ASVs within these water-wide ubiquitous sets were not necessarily absent from the biofilm, but did not reach full occupancy there (Fig 4b, Table S7). Overall, these patterns suggest a widespread component extending across water and biofilm, alongside taxa whose consistent occurrence was restricted to a subset of zones. Importantly, ubiquitous occurrence across both habitats did not correspond to similar abundance patterns (Fig. 4c). Among the ASVs ubiquitous across all four zones, most were abundant in the water (89.9–92.4% for bacteria and 80.0–86.7% for protists), whereas the proportion was lower in the biofilm (41.8% and 53.3%, respectively). Thus, the ubiquitous component shared between water and biofilm was predominantly abundant in the planktonic communities, whereas abundant and low-abundance taxa were more evenly represented in the biofilm (Fig 4C). This cross-habitat ubiquitous component included bacterial ASVs assigned to *hgcI clade* (Actinobacteriota), *Algoriphagus*, *Sediminibacterium* and NS9 marine group (Bacteroidota), and *Acidovorax* and *Limnohabitans* (Proteobacteria), as well as protist ASVs belonging to *Ceratium*, *Karlodinium* and *Parvodinium* (Alveolata), *Achnanthidium* and *Navicula* (Stramenopiles), and *Volvox* (Chlorophyta) (Table S8).

At the opposite end of the distribution, most ASVs detected exclusively in a given zone were characterized by restricted occupancy (Table S7). This was particularly evident in the biofilm, where 75.5% of non-ubiquitous biofilm-associated bacterial ASVs and 80.0% of their protist counterparts were low-abundance taxa restricted to a single sample (Table S7). In the individual water zones, unique ASVs were fewer and none were ubiquitous in either domain, with nearly all occurring at low abundance in only one to three samples. Thus, the large pool of ASVs associated to the biofilm primarily reflected taxa with restricted distributions rather than a larger habitat-specific ubiquitous component. In the water, ASVs detected exclusively in individual water zones also showed limited occupancy, whereas widespread ASVs were always shared beyond the individual water zone.

Having characterized the overall distribution of ASVs across habitats, we next examined the degree to which the abundance of ubiquitous taxa was consistent across the five lakes. Although ubiquitous ASVs were, by definition, detected in all five lakes of a given zone, their relative abundance between lakes in the given zone showed markedly different degrees of lake-to-lake consistency. Bacterial ubiquitous ASVs generally showed low to intermediate lake-to-lake variability: 31.9–37.1% had relatively consistent abundances (CV < 0.5), approximately half showed intermediate variability (46.9–52.6%; CV = 0.5–1), and only 12.9–16.9% showed high variability (CV > 1) (Table S9). Protist ASVs showed greater quantitative variation across lakes, with 36.9–49.6% of ubiquitous ASVs characterized by high variability and only 5.2–22.3% by low variability. This pattern was particularly evident among abundant ubiquitous protist ASVs, of which 42.3–65.8% showed high variability, compared with 9.1–26.2% of rare ubiquitous ASVs (Table S9). These proportions were broadly comparable across biofilm and water zones.

### Habitat-associated taxa identified by IndVal and Random Forest

To further investigate patterns of habitat association, we applied indicator value analysis (IndVal) and Random Forest classification (RF) to identify ASVs showing the strongest and most consistent associations with the sampled habitats. The two methods are complementary: IndVal identifies taxa with high fidelity and specificity to a habitat based on occurrence patterns, while RF captures taxa that discriminate habitats primarily through differences in relative abundance. Results from both approaches were subsequently related to the occupancy–abundance patterns described above.

Both methods consistently identified biofilm and water column as the primary axes of community differentiation, for both bacteria and protists. Habitat-associated taxa were predominantly linked either to the biofilm or to the combination of all three water zones — not to individual water zones — a pattern concordant with the occupancy analysis showing no zone-restricted ubiquitous ASVs in the water column.

### Bacteria

Among bacteria, IndVal identified 173 habitat-associated ASVs, of which 130 were exclusively associated with the biofilm (mean A = 0.95, B = 1), predominantly affiliated with Bacteroidota *(Flavobacterium, Cytophaga, Phaeodactylibacter*), Gemmatimonadota (*Gemmatimonas*) and Acidobacteriota (*Paludibaculum*). A further 39 ASVs were associated with the combination of all three water zones (mean A = 0.97, B = 1), including the *hgcI clade, CL500-29 marine group, Polynucleobacter, Polaromonas* and *Rhodoluna* (Table S10). The remaining 4 ASVs were associated with combinations of different two or three sampling zones (Table S10). No indicator ASVs were associated with individual water zones. RF selected 60 characteristic ASVs, of which 26 were characteristic of the biofilm, representing families such as Rhizobiaceae, Rhodobacteraceae, Saprospiraceae and Sphingomonadaceae, and 34 were characteristic of the water column, including the *hgcI clade, CL500-29 marine group* and typical freshwater genera such as *Limnohabitans, Rhodoluna, Polynucleobacter* and *Sediminibacterium* (Fig. 5a, Table S11). The slight asymmetry towards water-characteristic ASVs likely reflects the larger number of water samples available to the model (15 vs. 5 biofilm samples), which may improve detection of water-associated signal.

**Fig. 5.**
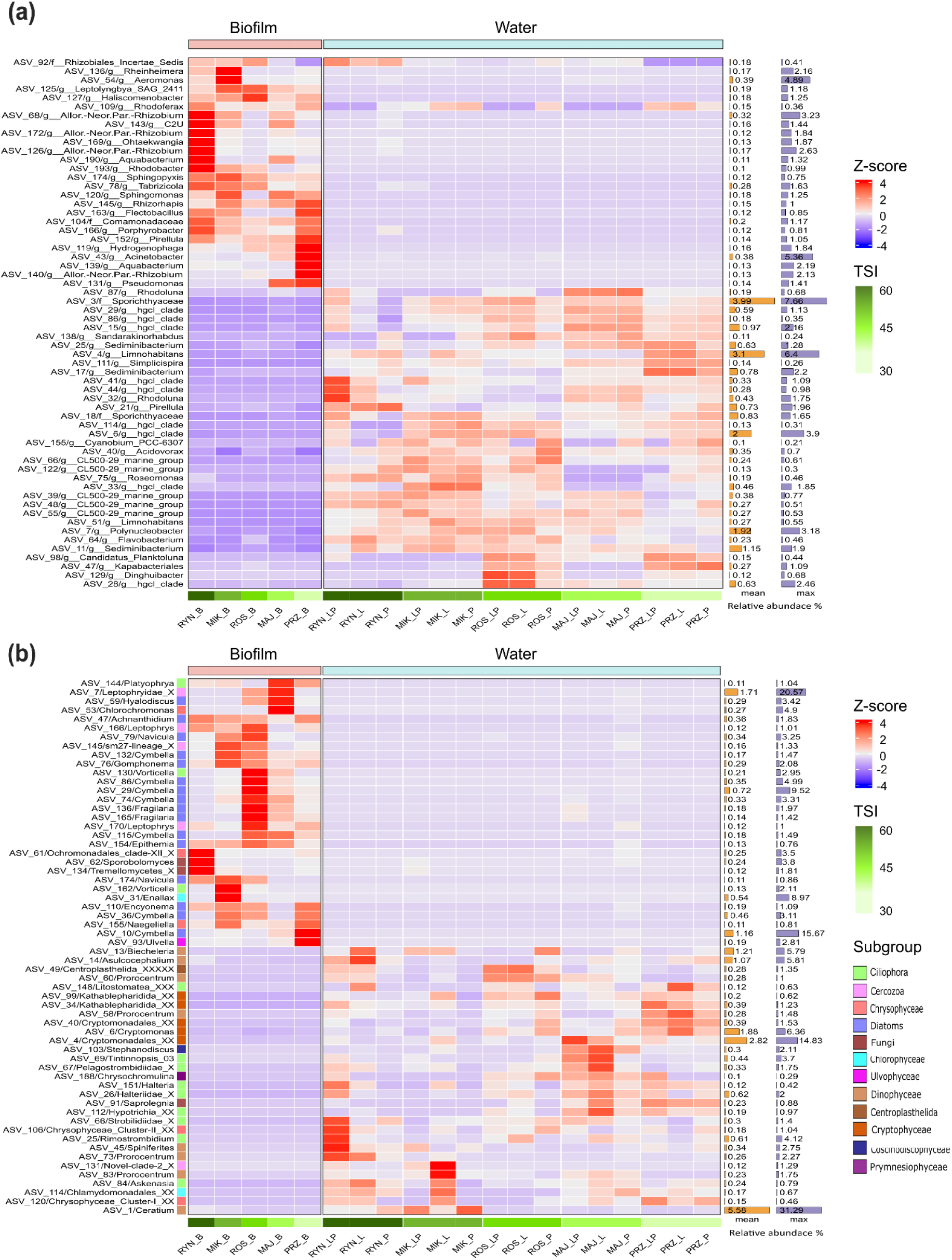
ASVs characteristic of biofilm and water communities identified by Random Forest (RF). Characteristic ASVs for each sample investigated were identified using a random forest algorithm. (**a**) Heatmap showing the abundance of the 60 ASVs selected by the random forest model across the 20 prokaryote samples. (**b**) Heatmap showing the abundance of the 60 ASVs selected by the model for protists. The first five samples represent biofilm communities, followed by water samples grouped by lake. Within each group, samples are arranged from higher to lower trophic level (left to right). The bar plots to the right of each heatmap display the overall relative abundance of each ASV across the entire dataset.

The two methods captured partially overlapping sets of ASVs (Table S12). Of the 60 top-ranking RF features, 26 were also IndVal indicators (Fig. 5, Table S12): 14 were enriched in biofilm (10 indicators of biofilm alone and 4 of biofilm combined with one or two water zones) and 12 were indicators of the three water zones combined. RF features occurred in at least three zones and discriminated habitats through shifts in relative abundance rather than occurrence. IndVal additionally captured habitat-restricted and rare ASVs largely missed by RF: none of the 43 biofilm-exclusive ASVs (36 rare, 7 abundant) was selected by RF, and only 2 of the 4 water-exclusive indicators, both abundant in all three water zones, were selected by RF. None of the rare indicators of biofilm (67 ASVs) or of the combined water zones (14 ASVs) was among the RF features (Table S12). Notably, IndVal identified four ASVs associated with the combination of all three water zones that were never detected in biofilm samples — a pattern not captured by occupancy analysis alone.

### Protists

For protists, both methods again identified biofilm and water as the primary axes of differentiation. IndVal identified 73 characteristic ASVs, with 51 associated exclusively with the biofilm (mean A = 0.94, B = 1), representing a taxonomically diverse assemblage including *Encyonopsis, Tetracystis, Salpingoecidae Group C2, Cladochytriales, Pleurostomatida* and *Hypotrichia*, and 20 associated with all three water zones (mean A = 0.98), including *Chrysophyceae Cluster I and II, Chrysochromulina, Spiniferites, Stephanodiscus, Tintinnopsis and Tintinnidium* (Table S10). The remaining two ASVs were associated with other sampling configurations: one with a combination of three zones and the other with the pelagic zone alone. RF identified 60 characteristic ASVs evenly distributed between biofilm and water column (30 ASVs each). Benthic diatoms were the most characteristic protist biofilm group, with 16 ASVs from nine genera — including *Achnanthidium, Epithemia and Leptophrys* — alongside Chlorophyta, Cercozoa, Fungi and Ciliophora, while the water-column assemblage included typical planktonic taxa such as *Halteria, Prorocentrum, Cryptomonas and Ceratium* (Fig. 5b, Table S11).

Agreement between methods was higher for water than for biofilm. RF selected 13 of the 20 water indicators, while only 9 of 51 biofilm indicators were among the RF features — largely because many biofilm-characteristic ASVs were abundant but patchy, captured by IndVal on occurrence grounds but not selected by RF on abundance grounds. This asymmetry further illustrates that biofilm community distinctiveness is characterised by both a consistent occurrence-based core and a broader set of abundance-variable habitat-associated taxa, whereas water-column differentiation is characterised more uniformly by abundant and widespread taxa.

Finally, the community-level physiological profiles and leucine-aminopeptidase activity data (Tables S2, S3 and Fig. S1) are consistent with those findings. Biofilm samples tended to show more similar metabolic profiles across lakes than planktonic samples, mirroring the greater compositional consistency observed for biofilm communities — particularly bacteria — in the analyses above.

## Discussion

Across the investigated lakes and sampling zones, biofilm and water communities showed some variation in community structure, despite being dominated by the same major phyla (Fig. 2a and c), consistent with previous studies reporting habitat-related differences in microbial composition [11, 10, 12]. Taxon distribution examination revealed a recurring positive abundance–occupancy relationship in both bacterial and protist assemblages across all the investigated zones (Fig. 3 and Table S7). Abundant ASVs, which represented a relatively small fraction of the detected taxa as also suggested by the Hill number profiles (Fig. 2b and Table S4), were predominantly associated with higher occupancy levels (Fig. 3 and Table S7). Conversely, the much larger low-abundance fraction was mainly composed of ASVs restricted to one or a few lakes (Figs. 2b and 3; Tables S4 and S7), consistent with abundance–occupancy relationships previously reported in freshwater microbial communities [38, 39]. This pattern may reflect dispersal and environmental filtering, although rare ubiquitous ASVs indicate that local rarity does not necessarily imply spatial restriction.

Beyond the general abundance–occupancy relationship, ubiquitous ASVs accounted for most of the relative abundance in all three aquatic zones for both bacteria and protists, while biofilms showed a greater contribution to total relative abundance from spatially restricted taxa (Table S7). This may partly reflect the microscale environmental heterogeneity typical of biofilms, which can provide diverse ecological niches favouring locally restricted taxa [40, 41], although in bacteria it may also be related to the larger number of ASVs detected in biofilms. Interestingly, the spatially restricted component distribution suggests contrasting abundance–occupancy patterns between bacteria and protists (Table S7), with some protist taxa reaching high local abundances despite limited spatial occurrence. This is consistent with the lake-to-lake patterns of ubiquitous taxa, with ubiquitous protists showing greater quantitative variability than bacteria, with their abundance more frequently concentrated in individual lakes (Tables S5 and S9). Thus, recurrent occurrence did not necessarily correspond to similar local abundance, potentially reflecting stronger influence of local environmental conditions and biotic interactions on protist abundance, allowing some taxa to reach high local abundances while remaining spatially restricted [42–44].

Habitat differentiation was also suggested in the distribution of shared and habitat-associated taxa. Biofilm harboured the largest pool of habitat-associated ASVs, particularly among bacteria, yet this pool was largely composed of low-abundance taxa detected in a single sample. Only 43 bacterial and 7 protist ASVs were consistently detected across all biofilm samples (Fig. 4; Table S7). The high ASV richness and apparent taxonomic distinctiveness of biofilm communities therefore largely reflected an extensive pool of rare and spatially restricted taxa, rather than a large, consistently occurring habitat-specific core (Table S7). However, these numbers should be interpreted with caution, as the limited number of sample per each zone may influence the identification of consistently detected ASVs, and additional sampling might alter these estimates. Such a pattern may reflect the stochastic colonisation of microhabitats within the biofilm matrix and the accumulation of taxa retained from the surrounding environment [12, 45, 46], potentially amplified by the higher bacterial richness of biofilms. Nevertheless, the existence of a small but consistent biofilm-associated component suggests that the biofilm environment may favour certain taxa across lakes. Importantly, intragenomic variation in rRNA gene sequences can generate multiple ASVs from a single species, potentially inflating apparent taxonomic turnover. Thus, taxon-level differences between biofilm communities may be smaller than ASV-level patterns suggest, although the observed restricted occupancy and low abundance remain relevant ecological patterns.

In contrast, no ASV was ubiquitous and simultaneously restricted to a single water zone: taxa widespread within a zone were always detected in the others (Fig. 4b; Table S7). This suggests a shared pool of widespread aquatic taxa, with zone-specific differences mainly involving the rare fraction, potentially reflecting water-column connectivity and mixing [47, 48, 12]. Many of this widespread planktonic pool were also ubiquitous in biofilms, although their relative abundances differed between habitats: shared ubiquitous ASVs were predominantly abundant in the water, whereas in the biofilm they were more evenly split between abundant and low-abundance taxa (Fig. 4c). This asymmetry is consistent with the idea that biofilm communities are partly assembled from the planktonic species pool, with water-borne taxa being continuously supplied to the biofilm but not necessarily thriving there [49, 50]. Thus, while many ASVs were detected across both habitats, their relative abundances differed between water and biofilm communities, a pattern consistent with habitat-associated environmental filtering reported in previous studies [51, 52].

While aquatic assemblages contained a substantial recurrent and abundant component shared across lakes and sampling zones, biofilms harboured an extensive pool of low-abundance taxa with restricted occupancy. Moreover, the pronounced variability observed among ubiquitous protist ASVs illustrates that recurring detection does not necessarily imply comparable quantitative importance across local assemblages. Across all analyses, bacteria and protists showed consistently contrasting patterns. Bacterial communities showed higher ASV richness in biofilm samples, a larger number of ASVs detected exclusively in the sampled biofilm communities, and relatively consistent abundance patterns across lakes. Protist ubiquitous ASVs, by contrast, showed greater quantitative variability across lakes regardless of habitat, suggesting that local conditions may play a more prominent role in protistan than in bacterial community structure.

For both bacteria and protists we further identified taxa most strongly associated with or informative of investigated habitats. IndVal revealed substantially more biofilm-associated than water-associated indicators, suggesting a recurrent habitat-associated component despite the high heterogeneity of biofilm communities (Table S10). Water indicators, in contrast, were predominantly associated with the three aquatic zones collectively rather than with individual zones (Table S10), supporting a shared planktonic component consistent with the occurrence patterns and bacterial PCoA described above (Figs. 2c and 4a). The recurrent component of aquatic assemblages therefore appeared to extend across the investigated sampling zones. This distribution may be compatible with microbial exchange within lakes, potentially facilitated by water movement and dispersal, although shared environmental conditions and the capacity of individual taxa to persist across different aquatic environments may also contribute [53, 50, 54].

Given the limitations of RF associated with the small sample size, its results were considered complementary to the IndVal analysis rather than as independent evidence of habitat differentiation. The partial overlap between IndVal and RF highlighted different aspects of ASV distribution across the sampled habitats (Table 12). While both approaches identified several ASVs in common, RF also selected broadly distributed ASVs that discriminated habitats mainly through differences in relative abundance, whereas IndVal identified additional rare or habitat-restricted ASVs (Fig. 5; Table S12).

Biofilm-associated bacterial and protistan ASVs included freshwater colonisers, primary producers and microbial consumers [49, 55]. Bacterial genera such as *Aeromonas*, *Acinetobacter* and *Flavobacterium* are associated with organic matter degradation and biofilm formation, while *Sphingomonas* and Comamonadaceae contribute to biofilm structure and carbon cycling [56–58]. Potential macrophyte associations were suggested by the *Allorhizobium–Neorhizobium–Pararhizobium–Rhizobium* complex, while several taxa, including *Sphingomonas*, *Hydrogenophaga*, *Pseudomonas*, *Rhodoferax* and *Flavobacterium*, are commonly associated with diatom-rich biofilms [59, 60]. Protistan community was characterised by phototrophic taxa, particularly benthic diatoms such as *Cymbella*, *Gomphonema*, *Navicula* and *Epithemia*, which contribute to biofilm structure and surface colonisation [3, 61], alongside fungal decomposers, chytrids and potential bacterial grazers such as *Vorticella* and *Aspidisca* [62–64]. Notably, *Epithemia* can harbour nitrogen-fixing cyanobacterial endosymbionts, potentially linking primary production with nitrogen input within freshwater biofilms [65].

Water-associated taxa were similarly representative of freshwater planktonic communities. Bacterial ASVs associated with water habitats according to IndVal, together with additional ASVs highlighted by the complementary RF analysis, included the hgcI clade (acI lineage), *Candidatus Planktoluna* (Luna/acIV), CL500-29, *Cyanobium_PCC-6307*, *Polynucleobacter* and *Rhodoluna*, belonging to lineages commonly reported in freshwater bacterioplankton, with hgcI and CL500-29 particularly associated with free-living communities [66]. *Cyanobium* is also frequently found in shallow and turbid lakes, where its small size may favour persistence under light limitation [67]. Similarly, protistan assemblages included typical planktonic taxa such as cryptophytes (*Cryptomonas*), kathablepharids, chrysophytes, the haptophyte *Chrysochromulina*, and planktonic ciliates including *Tintinnopsis lacustris*, *Tintinnidium*, oligotrichs and choreotrichs. Many of these taxa are mixotrophic or bacterivorous, potentially contributing to bacterioplankton grazing, while freshwater dinoflagellates such as *Ceratium hirundinella* and *Parvodinium inconspicuum* further reflected typical lacustrine plankton communities.

The taxonomic identity of shared ubiquitous ASVs provided additional insights into potential exchanges between habitats (Table S8). Many were abundant in water but rare on *Phragmites* surfaces, suggesting that planktonic taxa were largely passively retained within the biofilm matrix or the surrounding water film rather than being active biofilm members [68, 69]. Conversely, ASVs abundant in biofilms but rare in water included the Rhodobacteraceae *Rhodobacter* and *Tabrizicola* [70, 71] and the benthic diatoms *Gomphonema truncatum*, *Achnanthidium pyrenaicum* and *Navicula radiosa* [72]. Their detection in the water column, including the pelagic zone, is consistent with detachment from submerged surfaces and suggests that littoral biofilms may contribute, at least partially, to planktonic assemblages [73, 74].

Interestingly, the occurrence of certain ASVs exclusively in the sampled biofilm communities did not necessarily extend to higher taxonomic levels, as many genera and families containing these ASVs were also detected in water, often through different and less abundant ASVs. This may reflect fine-scale niche partitioning among closely related populations, including differences between particle-attached and free-living lifestyles [75], as well as the ability of some bacteria to switch between these states depending on resource availability [76]. For example, *Flavobacterium* and *Fluviicola* include both free-living and surface- or particle-associated populations [77], while Saprospiraceae are typically epiphytic or particle-associated polymer degraders and may enter the water through biofilm detachment [78, 79]. Similarly, env.OPS_17 includes non-motile bacteria growing on proteinaceous substrates [80], which may explain their association with biofilm-derived particles. Among protists, genera such as *Gomphonema* and *Navicula* include species and cryptic lineages with distinct ecological preferences, while peritrich ciliates and chytrids can alternate between sessile and dispersal stages [81, 82]. Thus, the presence of the same genus in both habitats does not necessarily indicate ecological equivalence among ASVs, and some variation may additionally reflect intragenomic rRNA gene heterogeneity, particularly in protists [83, 84]. These observations suggest that shared higher-level taxonomic identities do not necessarily imply equivalent ecological associations, highlighting the value of ASV-level resolution for detecting habitat-related patterns that may be obscured by broader taxonomic classifications.

### Limitations and future directions

Several limitations should be considered when interpreting these findings. First, biofilm communities were represented by a single composite sample per lake, precluding formal statistical comparison between habitat types and preventing assessment of within-lake spatial heterogeneity; future studies incorporating replicated biofilm samples would allow habitat effects to be distinguished from lake-level variation. Second, all samples were collected at a single time point — given that *Phragmites australis* biofilm communities change with season and biofilm developmental stage, the patterns described here represent a seasonal snapshot, and time-series sampling would allow successional and seasonal dynamics to be resolved. Third, the taxonomic resolution achievable with short amplicon reads is limited — many ASVs could only be assigned to family or genus level, and cryptic diversity, particularly among protists, may complicate ecological interpretation; long-read sequencing would allow more precise taxonomic assignment and better discrimination between genuine habitat specialists and population-level variants.

### Conclusions

Biofilm and planktonic microbial communities showed substantial taxonomic overlap, yet differed in the distribution and relative abundance of individual ASVs. Despite the large number of ASVs detected exclusively in *Phragmites australis* biofilms, most were rare and spatially restricted, while only a small fraction occurred consistently across all five lakes. In contrast, planktonic communities were characterised by a larger contribution of abundant and widespread ASVs, with substantial overlap among the three aquatic zones despite their different distances from reed stands.

Importantly, shared occurrence did not necessarily correspond to similar relative abundance across habitats or lakes. This distinction was particularly evident among protists, whose ubiquitous ASVs showed greater quantitative variability across lakes than bacterial ASVs, regardless of habitat. Bacterial communities, in turn, exhibited higher richness in biofilms and more consistent abundance patterns across lakes.

Together, these findings demonstrate that habitat-associated differentiation cannot be adequately described by taxon presence or absence alone. Integrating occurrence, relative abundance and spatial consistency provides a more nuanced understanding of microbial community organisation, revealing distinct patterns of habitat association in bacteria and protists within the same freshwater ecosystems.

## Acknowledgements

Freshwater sampling was conducted using the facilities of the KUMAK Masurian Centre for Biodiversity and Education in Urwitałt, Faculty of Biology, University of Warsaw. We would also like to thank all the MicroDivEr team members and Katarzyna Skłodowska who helped us with the sampling. We also thank Alessandro Cullichi for fruitful discussions and his critical feedback on the statistical interpretation of our results.

## Author contribution statement

Conceptualisation: AK, BK, VS. Developing methods: VS, MK, BK. Conducting the research: VS, MK, BK, AK. Data analysis: VS, BK. Data interpretation: VS, BK, AK. Preparation of figures and tables: VS. Writing original draft: VS. Writing review and editing: VS, AK, MK, BK.

## Data Availability Statement

The sequencing data have been deposited in the EMBL-EBI European Nucleotide Archive (ENA) under the Project ID PRJEB90292. All the rest of the Supplementary materials can be found in the Zenodo repository under DOI 10.5281/zenodo.17552625.

## References

1. Battin TJ, Kaplan LA, Denis Newbold J, Hansen CME (2003) Contributions of microbial biofilms to ecosystem processes in stream mesocosms. Nature 426:439–442. 10.1038/nature02152

2. Besemer K (2015) Biodiversity, community structure and function of biofilms in stream ecosystems. Research in Microbiology 166:774–781. 10.1016/j.resmic.2015.05.006

3. Battin TJ, Besemer K, Bengtsson MM, et al (2016) The ecology and biogeochemistry of stream biofilms. Nat Rev Microbiol 14:251–263. 10.1038/nrmicro.2016.15

4. Wijewardene L, Wu N, Fohrer N, Riis T (2022) Epiphytic biofilms in freshwater and interactions with macrophytes: Current understanding and future directions. Aquatic Botany 176:103467. 10.1016/j.aquabot.2021.103467

5. DeWolf EI, Calder WJ, Harrison JG, et al (2022) Aquatic Macrophytes Are Associated With Variation in Biogeochemistry and Bacterial Assemblages of Mountain Lakes. Frontiers in Microbiology Volume 12–2021: 10.3389/fmicb.2021.777084

6. Rusznyák A, Szabó G, Pollák B, et al (2007) Diversity of reed (Phragmites Australis) stem biofilm bacterial communities in two Hungarian soda lakes. Acta Microbiologica et Immunologica Hungarica 54:339–352. 10.1556/amicr.54.2007.4.2

7. Hiraki A, Tsuchiya Y, Fukuda Y, et al (2009) Analysis of How a Biofilm Forms on the Surface of the Aquatic Macrophyte Phragmites australis. Microb Environ 24:265–272. 10.1264/jsme2.ME09122

8. Tsuchiya Y, Hiraki A, Kiriyama C, et al (2011) Seasonal Change of Bacterial Community Structure in a Biofilm Formed on the Surface of the Aquatic Macrophyte Phragmites australis. Microb Environ 26:113–119. 10.1264/jsme2.ME10183

9. Rusznyák A, Vladár P, Molnár P, et al (2008) Cultivable bacterial composition and BIOLOG catabolic diversity of biofilm communities developed on Phragmites australis. Aquatic Botany 88:211–218. 10.1016/j.aquabot.2007.10.007

10. Souffreau C, Busschaert P, Denis C, et al (2018) A comparative hierarchical analysis of bacterioplankton and biofilm metacommunity structure in an interconnected pond system. Environmental Microbiology 20:1271–1282. 10.1111/1462-2920.14073

11. Parfenova VV, Gladkikh AS, Belykh OI (2013) Comparative analysis of biodiversity in the planktonic and biofilm bacterial communities in Lake Baikal. Microbiology 82:91–101. 10.1134/S0026261713010128

12. Robinson RL, Fisk AT, Crevecoeur S (2025) Temporal and Depth-Driven Variability of Pelagic Bacterial Communities in Lake Erie: Biofilm and Plankton Dynamics. Environ Microbiol Rep 17:e70079. 10.1111/1758-2229.70079

13. Siuda W, Grabowska K, Kaliński T, et al (2020) Trophic State, Eutrophication, and the Threats for Water Quality of the Great Mazurian Lake System. In: Korzeniewska E, Harnisz M (eds) Polish River Basins and Lakes – Part I. Springer International Publishing, Cham, pp 231–260

14. Carlson RE (1977) A trophic state index for lakes1. Limnology & Oceanography 22:361–369. 10.4319/lo.1977.22.2.0361

15. Kiersztyn B, Chróst R, Kaliński T, et al (2019) Structural and functional microbial diversity along a eutrophication gradient of interconnected lakes undergoing anthropopressure. Sci Rep 9:11144. 10.1038/s41598-019-47577-8

16. Chróst RJ, Siuda W (2006) Microbial production, utilization, and enzymatic degradation of organic matter in the upper trophogenic layer in the pelagial zone of lakes along a eutrophication gradient. Limnology & Oceanography 51:749–762. 10.4319/lo.2006.51.1_part_2.0749

17. Kiersztyn B, Siuda W, Chróst RJ (2012) Persistence of bacterial proteolytic enzymes in lake ecosystems. FEMS Microbiol Ecol 80:124–134. 10.1111/j.1574-6941.2011.01276.x

18. Romac S (2022) Eukaryotes 18S-V4 rRNA Metabarcoding PCR protocol for NGS Illumina sequencing. protocols.io

19. Romac S (2022) Prokaryotes 16S-V4V5 rRNA Metabarcoding PCR protocol for NGS Illumina sequencing. protocols.io. 10.17504/protocols.io.bzwwp7fe

20. Andrews (2010) A Quality Control Tool for High Throughput Sequence Data [Online]. Available online at. http://www.bioinformatics.babraham.ac.uk/projects/fastqc/

21. Bolyen, Rideout, Dillon, et al. (2019) Reproducible, interactive, scalable and extensible microbiome data science using QIIME 2. Nat Biotechnol 37:852–857. 10.1038/s41587-019-0209-9

22. Martin (2011) Cutadapt removes adapter sequences from highthroughput sequencing reads. EMBnet J 17:10. 10.14806/ej.17.1.200

23. Callahan BJ, McMurdie PJ, Rosen MJ, et al (2016) DADA2: High-resolution sample inference from Illumina amplicon data. Nat Methods 13:581–583. 10.1038/nmeth.3869

24. Rognes T, Flouri T, Nichols B, et al (2016) VSEARCH: a versatile open source tool for metagenomics. PeerJ 4:e2584. 10.7717/peerj.2584

25. Quast C, Pruesse E, Yilmaz P, et al (2012) The SILVA ribosomal RNA gene database project: improved data processing and web-based tools. Nucleic Acids Research 41:D590–D596. 10.1093/nar/gks1219

26. Guillou L, Bachar D, Audic S, et al (2012) The Protist Ribosomal Reference database (PR2): a catalog of unicellular eukaryote Small Sub-Unit rRNA sequences with curated taxonomy. Nucleic Acids Research 41:D597–D604. 10.1093/nar/gks1160

27. Allaire (2012) RStudio: integrated development environment for R. Boston: The R Project for Statistical Computing Vol. 770:165–71

28. McMurdie PJ, Holmes S (2013) phyloseq: An R Package for Reproducible Interactive Analysis and Graphics of Microbiome Census Data. PLoS ONE 8:e61217. 10.1371/journal.pone.0061217

29. Karlicki M, Bednarska A, Hałakuc P, et al (2024) Spatio-temporal changes of small protist and free-living bacterial communities in a temperate dimictic lake: insights from metabarcoding and machine learning. FEMS Microbiology Ecology 100:fiae104. 10.1093/femsec/fiae104

30. Barnett, Arts, Penders (2021) microViz: an R package for microbiome data visualization and statistics. Journal of Open Source Software, 6(63), 3201, 10.21105/joss.03201. Journal of Open Source Software 6(63):3201. 10.21105/joss.03201

31. Lahti L, Shetty S (2017) Microbiome R package. Boston: Bioconductor. 10.18129/B9.bioc.microbiome

32. Hsieh TC, Ma KH, Chao A (2016) iNEXT: an R package for rarefaction and extrapolation of species diversity ( H ill numbers). Methods Ecol Evol 7:1451–1456. 10.1111/2041-210X.12613

33. Crump BC, Amaral-Zettler LA, Kling GW (2012) Microbial diversity in arctic freshwaters is structured by inoculation of microbes from soils. The ISME Journal 6:1629–1639. 10.1038/ismej.2012.9

34. Peter H, Jeppesen E, De Meester L, Sommaruga R (2018) Changes in bacterioplankton community structure during early lake ontogeny resulting from the retreat of the Greenland Ice Sheet. The ISME Journal 12:544–555. 10.1038/ismej.2017.191

35. Li C, Jin L, Zhang C, et al (2023) Destabilized microbial networks with distinct performances of abundant and rare biospheres in maintaining networks under increasing salinity stress. iMeta 2:e79. 10.1002/imt2.79

36. Cáceres MD, Legendre P (2009) Associations between species and groups of sites: indices and statistical inference. Ecology 90:3566–3574. 10.1890/08-1823.1

37. Breiman L (2001) Random Forests. Machine Learning 45:5–32. 10.1023/A:1010933404324

38. Liu L, Yang J, Yu Z, Wilkinson DM (2015) The biogeography of abundant and rare bacterioplankton in the lakes and reservoirs of China. The ISME Journal 9:2068–2077. 10.1038/ismej.2015.29

39. Niño-García JP, Ruiz-González C, Del Giorgio PA (2016) Landscape-scale spatial abundance distributions discriminate core from random components of boreal lake bacterioplankton. Ecology Letters 19:1506–1515. 10.1111/ele.12704

40. Flemming H-C, Wingender J (2010) The biofilm matrix. Nat Rev Microbiol 8:623–633. 10.1038/nrmicro2415

41. Jo J, Price-Whelan A, Dietrich LEP (2022) Gradients and consequences of heterogeneity in biofilms. Nature Reviews Microbiology 20:593–607. 10.1038/s41579-022-00692-2

42. Simon M, López-García P, Deschamps P, et al (2015) Marked seasonality and high spatial variability of protist communities in shallow freshwater systems. The ISME Journal 9:1941–1953. 10.1038/ismej.2015.6

43. Bock C, Jensen M, Forster D, et al (2020) Factors shaping community patterns of protists and bacteria on a European scale. Environmental Microbiology 22:2243–2260. 10.1111/1462-2920.14992

44. Liu K, Liu Y, Hu A, et al (2020) Different community assembly mechanisms underlie similar biogeography of bacteria and microeukaryotes in Tibetan lakes. FEMS Microbiology Ecology 96:fiaa071. 10.1093/femsec/fiaa071

45. Ezzat L, Fodelianakis S, Kohler TJ, et al (2022) Benthic Biofilms in Glacier-Fed Streams from Scandinavia to the Himalayas Host Distinct Bacterial Communities Compared with the Streamwater. Applied and Environmental Microbiology 88:e00421–22. 10.1128/aem.00421-22

46. Flemming H-C, van Hullebusch ED, Neu TR, et al (2023) The biofilm matrix: multitasking in a shared space. Nature Reviews Microbiology 21:70–86. 10.1038/s41579-022-00791-0

47. Shade A, Jones SE, McMahon KD (2008) The influence of habitat heterogeneity on freshwater bacterial community composition and dynamics. Environmental Microbiology 10:1057–1067. 10.1111/j.1462-2920.2007.01527.x

48. Shade A, Chiu C-Y., McMahon KD (2010) Differential bacterial dynamics promote emergent community robustness to lake mixing: an epilimnion to hypolimnion transplant experiment. Environmental Microbiology 12:455–466. 10.1111/j.1462-2920.2009.02087.x

49. Besemer K, Peter H, Logue JB, et al (2012) Unraveling assembly of stream biofilm communities. The ISME Journal 6:1459–1468. 10.1038/ismej.2011.205

50. Lindström ES, Langenheder S (2012) Local and regional factors influencing bacterial community assembly. Environ Microbiol Rep 4:1–9. 10.1111/j.1758-2229.2011.00257.x

51. Zhu C, Bass D, Wang Y, et al (2020) Environmental Parameters and Substrate Type Drive Microeukaryotic Community Structure During Short-Term Experimental Colonization in Subtropical Eutrophic Freshwaters. Front Microbiol 11:555795. 10.3389/fmicb.2020.555795

52. Gweon HS, Bowes MJ, Moorhouse HL, et al (2021) Contrasting community assembly processes structure lotic bacteria metacommunities along the river continuum. Environmental Microbiology 23:484–498. 10.1111/1462-2920.15337

53. Pandit SN, Kolasa J, Cottenie K (2009) Contrasts between habitat generalists and specialists: an empirical extension to the basic metacommunity framework. Ecology 90:2253–2262. 10.1890/08-0851.1

54. Székely AJ, Langenheder S (2014) The importance of species sorting differs between habitat generalists and specialists in bacterial communities. FEMS Microbiol Ecol 87:102–112. 10.1111/1574-6941.12195

55. Battin TJ, Sloan WT, Kjelleberg S, et al (2007) Microbial landscapes: new paths to biofilm research. Nat Rev Microbiol 5:76–81. 10.1038/nrmicro1556

56. Czieborowski M, Hübenthal A, Poehlein A, et al (2020) Genetic and physiological analysis of biofilm formation on different plastic surfaces by Sphingomonas sp. strain S2M10 reveals an essential function of sphingan biosynthesis. Microbiology 166:918–935. 10.1099/mic.0.000961

57. Grilo ML, Pereira A, Sousa-Santos C, et al (2021) Climatic Alterations Influence Bacterial Growth, Biofilm Production and Antimicrobial Resistance Profiles in Aeromonas spp. Antibiotics 10:1008. 10.3390/antibiotics10081008

58. Pandolfo E, Barra Caracciolo A, Rolando L (2023) Recent Advances in Bacterial Degradation of Hydrocarbons. Water 15:375. 10.3390/w15020375

59. Grossart, Levold F, Allgaier M, et al (2005) Marine diatom species harbour distinct bacterial communities. Environmental Microbiology 7:860–873. 10.1111/j.1462-2920.2005.00759.x

60. Bruckner CG, Bahulikar R, Rahalkar M, et al (2008) Bacteria Associated with Benthic Diatoms from Lake Constance: Phylogeny and Influences on Diatom Growth and Secretion of Extracellular Polymeric Substances. Appl Environ Microbiol 74:7740–7749. 10.1128/AEM.01399-08

61. Fernández HR, Gonzalez Achem AL, Correa M, Albarracín VH (2019) Microbialite-like structures in Cladophora sp. (Ulvophyceae) mats from a subtropical Andean basin: ecological implications. RBT 67:. 10.15517/rbt.v67i1.31542

62. Grossart, Wurzbacher C, James TY, Kagami M (2016) Discovery of dark matter fungi in aquatic ecosystems demands a reappraisal of the phylogeny and ecology of zoosporic fungi. Fungal Ecology 19:28–38. 10.1016/j.funeco.2015.06.004

63. Roberts C, Allen R, Bird KE, Cunliffe M (2020) Chytrid fungi shape bacterial communities on model particulate organic matter. Biol Lett 16:20200368. 10.1098/rsbl.2020.0368

64. Klawonn I, Van Den Wyngaert S, Iversen MH, et al (2023) Fungal parasitism on diatoms alters formation and bio–physical properties of sinking aggregates. Commun Biol 6:206. 10.1038/s42003-023-04453-6

65. Moulin SLY, Frail S, Braukmann T, et al (2024) The endosymbiont of *Epithemia clementina* is specialized for nitrogen fixation within a photosynthetic eukaryote. ISME Communications 4:ycae055. 10.1093/ismeco/ycae055

66. Newton RJ, Jones SE, Eiler A, et al (2011) A Guide to the Natural History of Freshwater Lake Bacteria. Microbiol Mol Biol Rev 75:14–49. 10.1128/MMBR.00028-10

67. Bernát G, Zavřel T, Kotabová E, et al (2021) Photomorphogenesis in the Picocyanobacterium Cyanobium gracile Includes Increased Phycobilisome Abundance Under Blue Light, Phycobilisome Decoupling Under Near Far-Red Light, and Wavelength-Specific Photoprotective Strategies. Front Plant Sci 12:612302. 10.3389/fpls.2021.612302

68. Al-hadithi SA, Goulder R (1989) Physiological state of epiphytic bacteria on submerged stems of the reed Phragmites australis compared with planktonic bacteria in gravel-pit ponds. Journal of Applied Bacteriology 66:107–117. 10.1111/j.1365-2672.1989.tb02460.x

69. Yan D, Xia P, Song X, et al (2019) Community structure and functional diversity of epiphytic bacteria and planktonic bacteria on submerged macrophytes in Caohai Lake, southwest of China. Ann Microbiol 69:933–944. 10.1007/s13213-019-01485-4

70. Di Pippo F, Venezia C, Sighicelli M, et al (2020) Microplastic-associated biofilms in lentic Italian ecosystems. Water Research 187:116429. 10.1016/j.watres.2020.116429

71. Sheu C, Li Z-H, Sheu S-Y, et al (2020) Tabrizicola oligotrophica sp. nov. and Rhodobacter tardus sp. nov., two new species of bacteria belonging to the family Rhodobacteraceae. International Journal of Systematic and Evolutionary Microbiology 70:6266–6283. 10.1099/ijsem.0.004526

72. Nemes-Kókai Z, Borics G, Csépes E, et al (2024) Role of microhabitats in shaping diversity of periphytic diatom assemblages. Hydrobiologia 851:959–972. 10.1007/s10750-023-05336-x

73. Istvànovic V, Honti M (2011) Phytoplankton growth in three rivers: The role of meroplankton and the benthic retention hypothesis. Limnology & Oceanography 56:1439–1452. 10.4319/lo.2011.56.4.1439

74. Jin H, Van Leeuwen CHA, Van De Waal DB, Bakker ES (2022) Impacts of sediment resuspension on phytoplankton biomass production and trophic transfer: Implications for shallow lake restoration. Science of The Total Environment 808:152156. 10.1016/j.scitotenv.2021.152156

75. Hunt DE, David LA, Gevers D, et al (2008) Resource Partitioning and Sympatric Differentiation Among Closely Related Bacterioplankton. Science 320:1081–1085. 10.1126/science.1157890

76. Grossart H (2010) Ecological consequences of bacterioplankton lifestyles: changes in concepts are needed. Environ Microbiol Rep 2:706–714. 10.1111/j.1758-2229.2010.00179.x

77. Kirchman D (2002) The ecology of Cytophaga–Flavobacteria in aquatic environments. FEMS Microbiology Ecology 39:91–100. 10.1016/S0168-6496(01)00206-9

78. McIlroy SJ, Nielsen PH (2014) The Family Saprospiraceae. In: Rosenberg E, DeLong EF, Lory S, et al (eds) The Prokaryotes. Springer Berlin Heidelberg, Berlin, Heidelberg, pp 863–889

79. Bižić-Ionescu M, Ionescu D, Grossart H-P (2018) Organic Particles: Heterogeneous Hubs for Microbial Interactions in Aquatic Ecosystems. Front Microbiol 9:2569. 10.3389/fmicb.2018.02569

80. Vieira S, Riedel T, Geppert A, et al (2023) Aurantibacillus circumpalustris gen. nov., sp. nov., the first characterized representative of the Bacteroidota candidate family env.OPS 17 and proposal of Aurantibacillaceae fam. nov. International Journal of Systematic and Evolutionary Microbiology 73:. 10.1099/ijsem.0.006134

81. Abarca N, Jahn R, Zimmermann J, Enke N (2014) Does the Cosmopolitan Diatom Gomphonema parvulum (Kützing) Kützing Have a Biogeography? PLoS ONE 9:e86885. 10.1371/journal.pone.0086885

82. Kagami M, De Bruin A, Ibelings BW, Van Donk E (2007) Parasitic chytrids: their effects on phytoplankton communities and food-web dynamics. Hydrobiologia 578:113–129. 10.1007/s10750-006-0438-z

83. Gong J, Dong J, Liu X, Massana R (2013) Extremely High Copy Numbers and Polymorphisms of the rDNA Operon Estimated from Single Cell Analysis of Oligotrich and Peritrich Ciliates. Protist 164:369–379. 10.1016/j.protis.2012.11.006

84. Trubovitz S, Sandin MM, Caron DA (2025) Genetic variability in microbial eukaryotes reshapes marine biodiversity assessment in the age of amplicon sequencing. PLoS One 20:e0326053. 10.1371/journal.pone.0326053

