## Supplementary Figures for "Diversity and habitat-associated distribution of bacterial and protistan communities in *Phragmites australis* biofilms and surrounding waters of Masurian lakes (Poland)"

**Supplementary Tables**

Supplementary Tables are available in "Supplementary_Tables.xlsx". The file contains the following tables:

**Table S1.** Metadata table for each sample including sample abbreviation, lake name, matrix type, sampling zone, sampling date, basic physico-chemical parameters, and TSI values.

**Table S2.** Functional physiological profiling of biofilm and water samples based on the Biolog EcoPlate™ assay. The left section of the table reports the percentage of metabolized carbon sources for each environmental sample; the right section presents the median utilization values of substrates grouped by chemical class (e.g., carbohydrates, amino acids, etc.)

**Table S3.** Leucine-aminopeptidase activity and kinetic parameters. For biofilms, Vmax values are normalized using volume coefficients based on the mean volume of Phragmites australis stems (see box on the right)

**Table S4.** Hill numbers (q0, q1, and q2) for bacterial and protist communities across samples. Sample codes indicate lake and sampling zone: B, biofilm; LI, littoral water inside the Phragmites australis bed; LP, littoral water outside the P. australis bed; P, pelagic water. The dataset column indicates whether numbers refer to bacterial (16S) or protist (18S) data

**Table S5.** ASVs with max_fraction > 0.5 (a single lake explains over 50% of total abundance)

**Table S6.** Percentage of raw reads that passed denoising and filtering steps during DADA2 processing in QIIME2 and percentage of ASVs before and after taxa filtering for the 16S and 18S datasets

**Table S7.** Abundance–occupancy distribution and habitat specificity of bacterial and protist ASVs across sampling zones. ASVs were classified according to their occupancy across the five investigated lakes (from 1/5 to 5/5) and their relative abundance as abundant (>0.1%) or rare (<0.1%). For each sampling zone and domain, the table reports the number and proportion of abundant and rare ASVs within each occupancy category, their contribution to total relative abundance, and the number of ASVs unique to each sampling zone. ASVs occurring in 5/5 lakes were considered ubiquitous. Unique ASVs were defined as ASVs detected exclusively in a given sampling zone and absent from the other three zones

**Table S8.** The table includes ASVs that were ubiquitous and unique to the Phragmites australis surface biofilm, as well as ubiquitous ASVs shared among all four habitats (Phragmites surface biofilm, littoral water inside the Phragmites australis field, littoral water outside the Phragmites australis field, and pelagic water). For each ASV, taxonomic affiliation and abundance–occupancy classification are reported

**Table S9.** CV distribution per zone, split by category (Abundant & ubiquitous vs Rare & ubiquitous)

**Table S10.** Indicator ASVs identified by IndVal analysis for bacterial (16S rRNA gene) and eukaryotic (18S rRNA gene) communities across the four sampled zones. For each ASV, the table reports the zone or combination of zones for which it serves as an indicator, the specificity (A) and fidelity (B) components, the IndVal statistic (stat), the associated p-value, zone-specific specificity values (s.), and the full taxonomic classification from domain to species level

**Table S11.** List of ASVs characteristic of water and biofilm, selected using the Random Forest (RF) model, with their corresponding genus-level assignments. When genus-level identification was not available, the last taxonomic level assigned was used for classification. Prokaryotic ASVs are shown on the left, and protist ASVs on the right. ASVs typical of biofilm are highlighted in green (prokaryotes) and yellow (protists). ASVs most characteristic of water are not highlighted

**Table S12.** Comparison of ASVs identified by Random Forest with Indicators for 16S and 18S. The table shows in how many zones the same ASV occurs and if it's also identified as an indicator

### Supplementary Figures

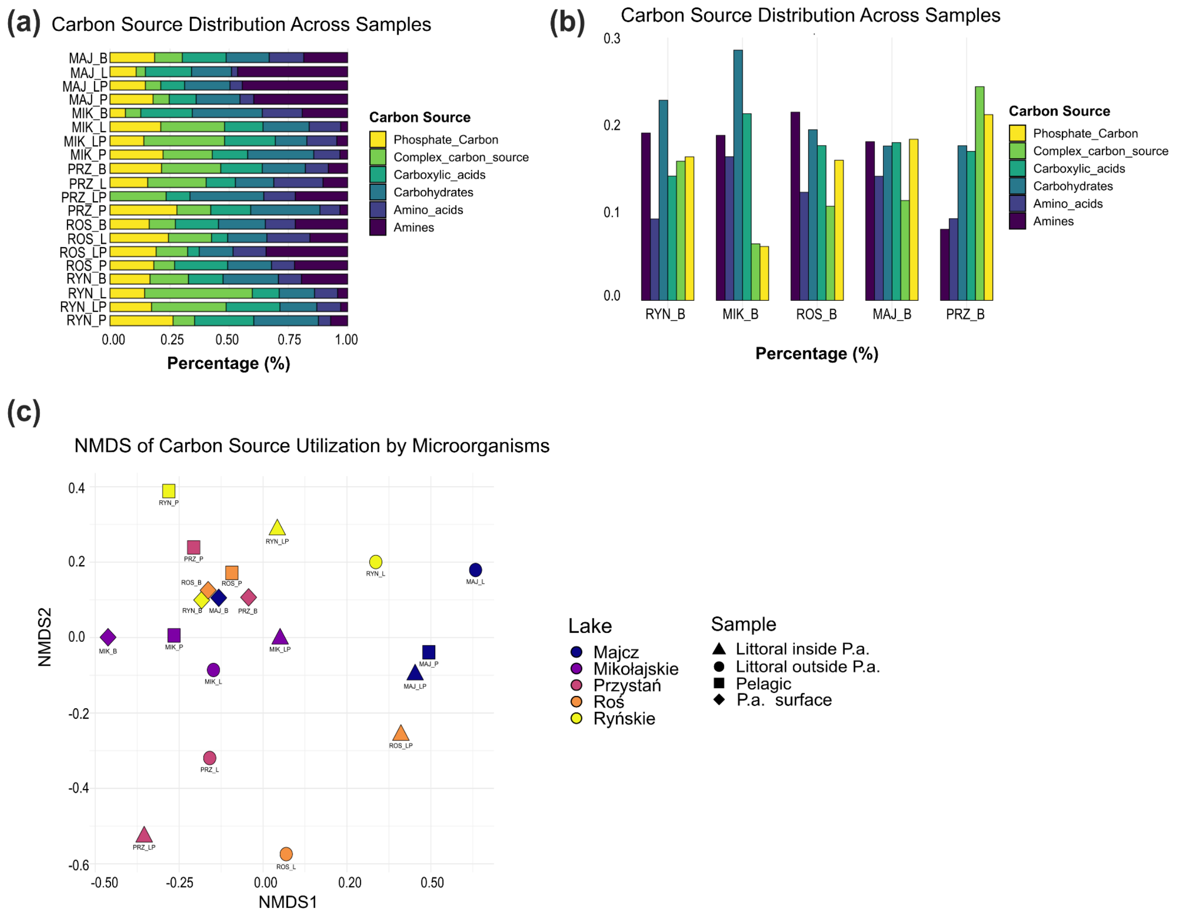


**Fig. 1** Community-level physiological profiles of biofilm and planktonic communities assessed using Biolog EcoPlates. Biofilm samples showed broadly similar metabolic profiles across lakes while planktonic profiles were more variable, consistent with the compositional patterns described in the main text. Results represent single samples per lake and were not normalised to biomass; therefore, they should be interpreted as indicative only. (**a**) Average relative utilisation of carbon sources by compound type. (**b**) Carbon source utilisation among biofilm samples. (**c**) NMDS ordination of Bray–Curtis dissimilarities across all samples; used for descriptive purposes only, no statistical inference drawn

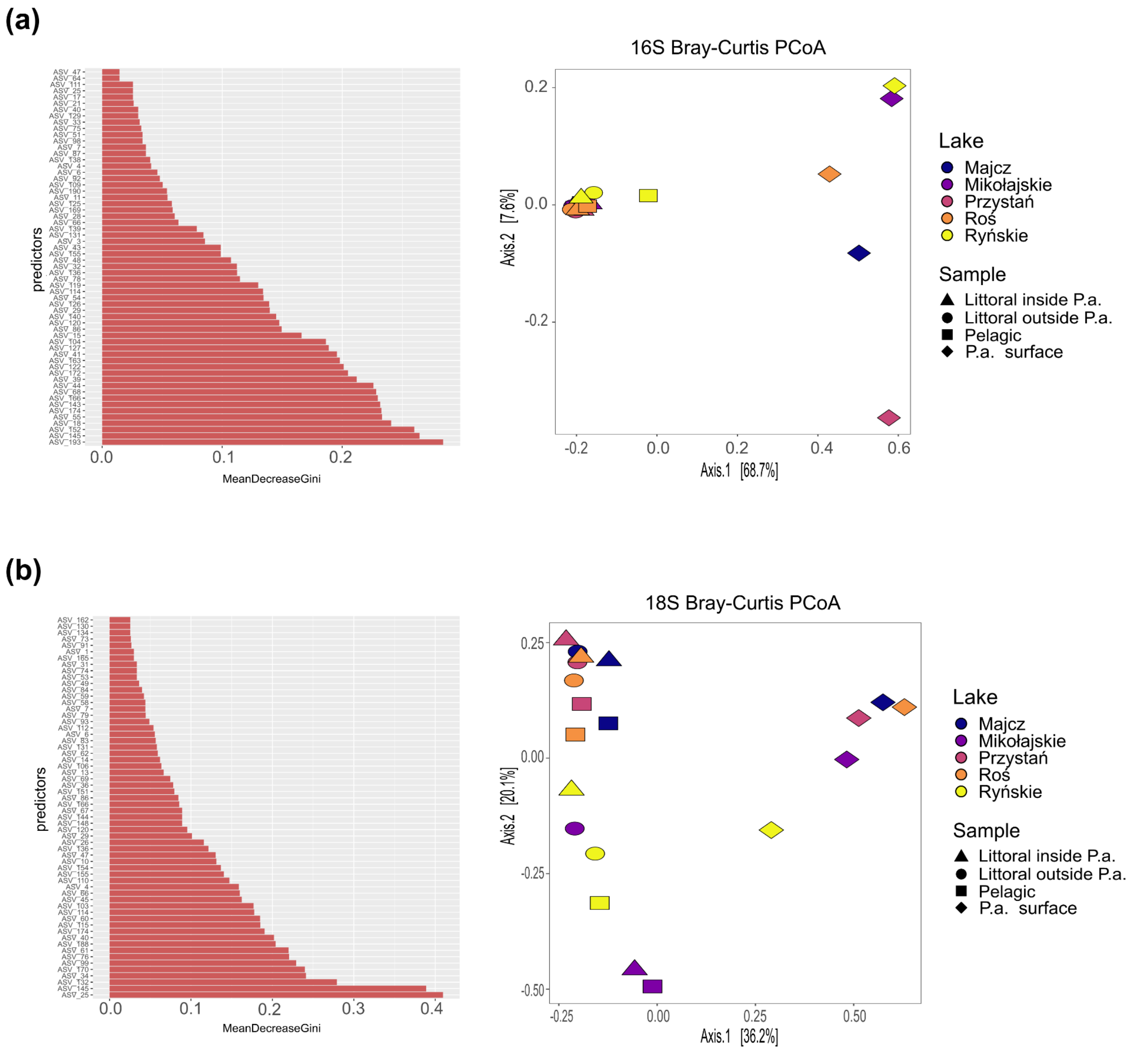


**Fig. S2** Mean Gini index and ordination of key ASVs differentiating water and biofilm samples for (**a**) prokaryotes and (**b**) protists. Left panel: Mean Gini index of the first 60 predictors selected by the Random Forest model. Right panel: Bray-Curtis PCoA ordination plots based on ASVs selected by Random Forest models. Shapes represent the sample type, and colours indicate different lakes

(a)

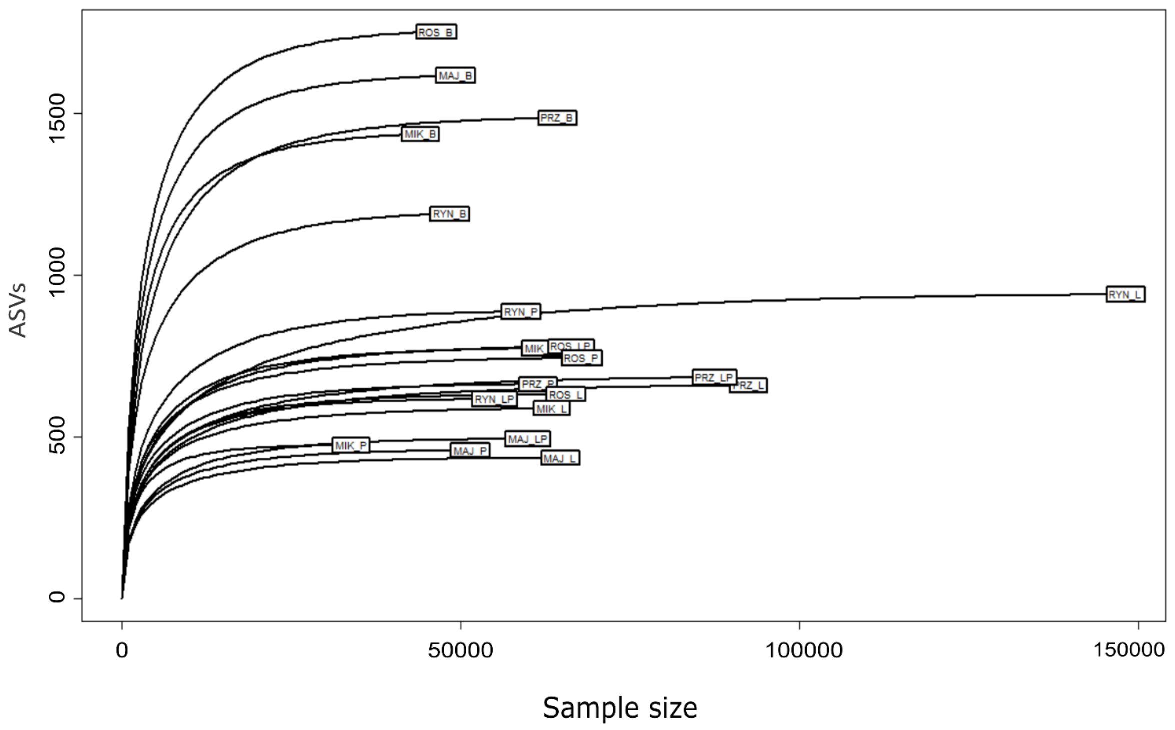


(b)

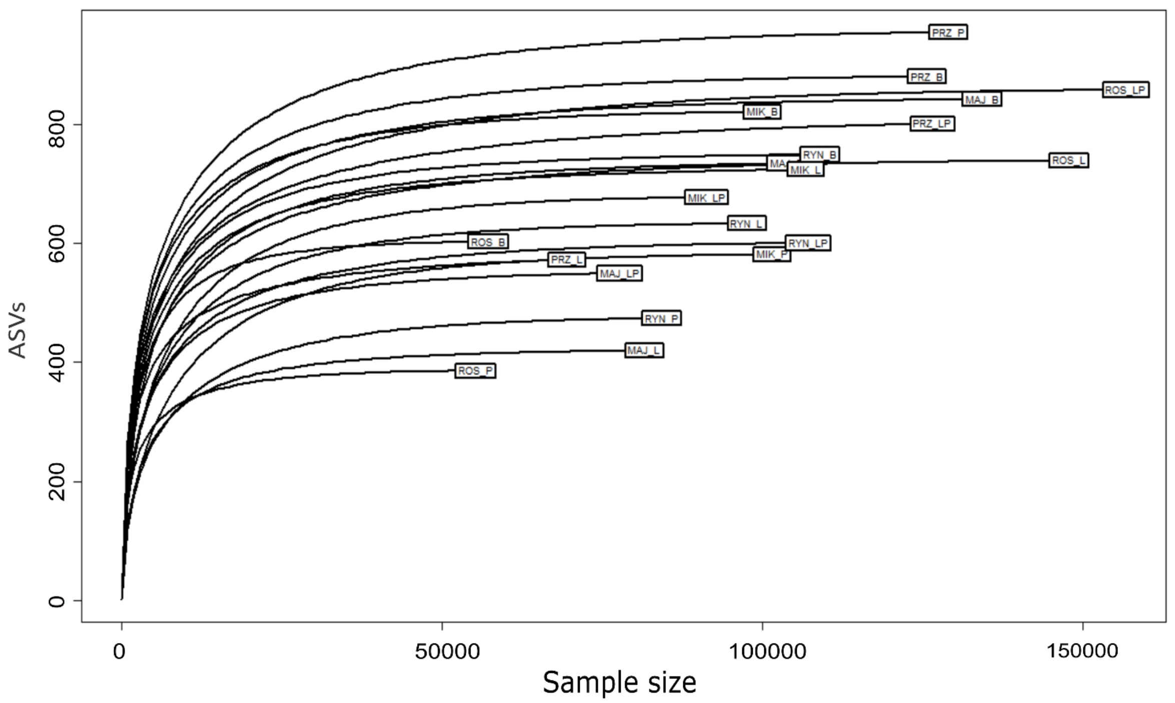


**Fig. S3** Rarefaction Curves. (a) The 16S rarefaction curve, and (b) the 18S curve

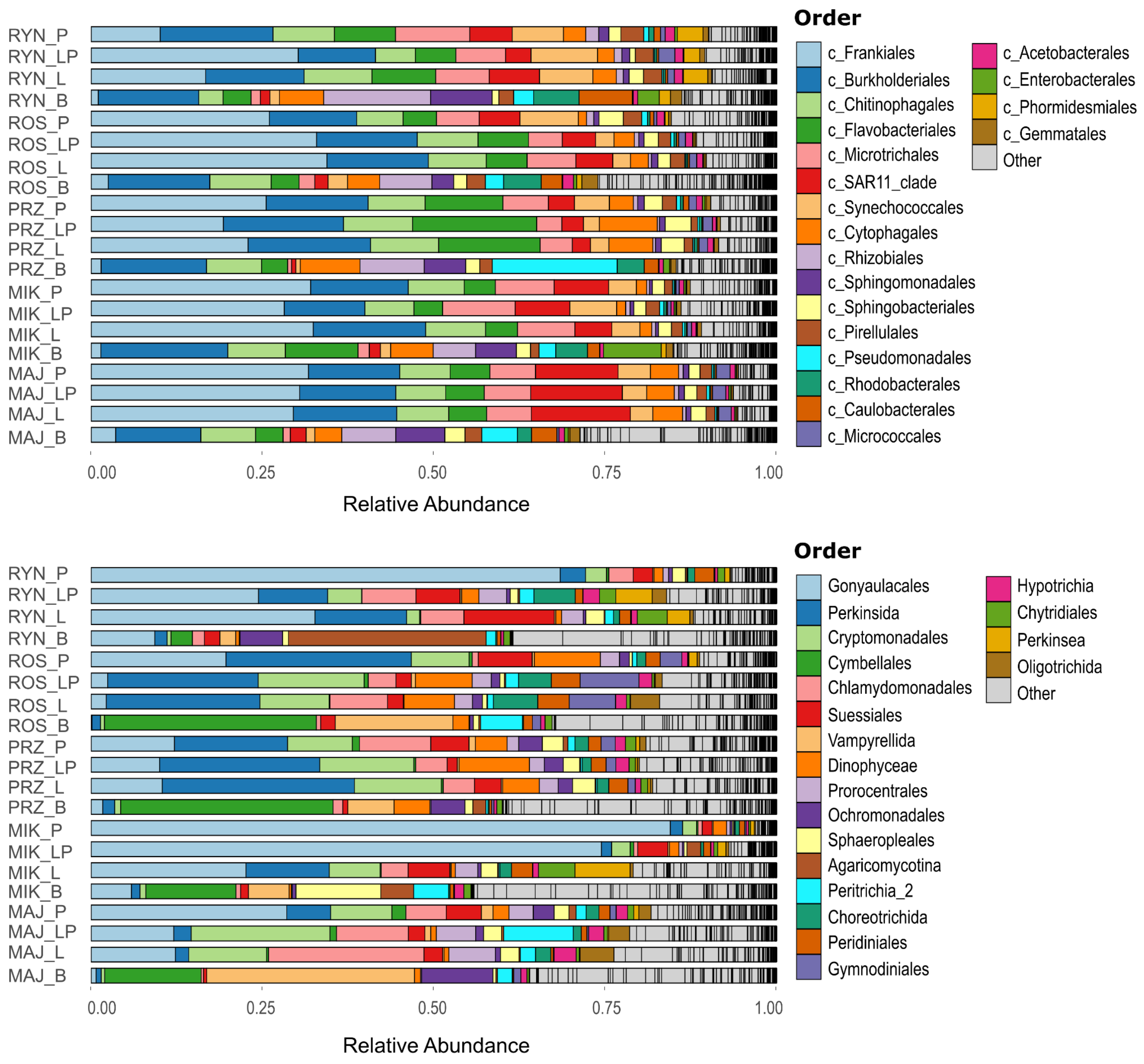


**Fig. S4** Taxonomic composition of the bacterial communities. The bar plot shows the relative abundances of the most abundant bacterial (upper panel) and protist (lower panel) taxa at the order level across biofilm and water samples from the five studied lakes. All remaining taxa are grouped as “Other”.
